# SAP: Synteny-aware gene function prediction for bacteria using protein embeddings

**DOI:** 10.1101/2023.05.02.539034

**Authors:** Aysun Urhan, Bianca-Maria Cosma, Ashlee M. Earl, Abigail L. Manson, Thomas Abeel

## Abstract

**Motivation:** Today, we know the function of only a small fraction of the protein sequences predicted from genomic data. This problem is even more salient for bacteria, which represent some of the most phylogenetically and metabolically diverse taxa on Earth. This low rate of bacterial gene annotation is compounded by the fact that most function prediction algorithms have focused on eukaryotes, and conventional annotation approaches rely on the presence of similar sequences in existing databases. However, often there are no such sequences for novel bacterial proteins. Thus, we need improved gene function prediction methods tailored for prokaryotes. Recently, transformer-based language models - adopted from the natural language processing field - have been used to obtain new representations of proteins, to replace amino acid sequences. These representations, referred to as protein embeddings, have shown promise for improving annotation of eukaryotes, but there have been only limited applications on bacterial genomes.

**Results:** To predict gene functions in bacteria, we developed SAP, a novel synteny-aware gene function prediction tool based on protein embeddings from state-of-the-art protein language models. SAP also leverages the unique operon structure of bacteria through conserved synteny. SAP outperformed both conventional sequence-based annotation methods and state-of-the-art methods on multiple bacterial species, including for distant homolog detection, where the sequence similarity to the proteins in the training set was as low as 40%. Using SAP to identify gene functions across diverse enterococci, of which some species are major clinical threats, we identified 11 previously unrecognized putative novel toxins, with potential significance to human and animal health.

**Availability:** https://github.com/AbeelLab/sap

**Supplementary information:** Supplementary data are available at *Bioinformatics* online.

## 1 Introduction

With increasing volumes of sequencing data from high-throughput technologies, the observed diversity of protein sequences is increasing faster than our knowledge of its functional significance. Given costs and the inability to scale experimental and other manual approaches for function prediction, computational approaches have a critical role in deciphering functional diversity. Most state-of-the-art protein function prediction methods have focused on annotation of eukaryotic proteins, leaving a gap in our understanding of the vast landscape of protein diversity among the bacterial domain, representing some of the most phylogenetically and metabolically diverse taxa.

Conventional approaches to predicting gene function rely on sequence homology. Initial methods employed sequence search tools such as BLAST or DIAMOND to query a database of known protein sequences and their functions (Altschul *et al*., 1990; Buchfink *et al*., 2021). While useful, these methods are limited by the completeness (i.e., lack of homologs) and fidelity (i.e., inclusion of annotation errors) of the databases they use. Furthermore, it is often difficult to determine a proper threshold to transfer gene function, resulting in low sensitivity and specificity (Zhou *et al*., 2019). With increasing volumes of data, machine learning techniques for function prediction have been explored, including using features derived from the sequence of interest in supervised machine learning models, such as multilayer perceptrons, support vector machines, and k-nearest neighbor (knn) algorithms (Jensen *et al*., 2003; Törönen *et al*., 2018). In the most recent Critical Assessment of Functional Annotation (CAFA), a challenge established to evaluate the state-of-the-art in automated function prediction, GOLabeler was the top performing method for predicting molecular function ontologies Zhou *et al*. (2019) (Zhou et al. 2019) by integrating sequence alignments, domain and motif information, and biophysical properties of protein sequences to predict gene function (You *et al*., 2018).

More recently, deep learning methods that leverage ideas from natural language processing (NLP) have gained attention for gene function prediction. Deep learning-based protein language models were recently used to extract embedding vectors for protein sequences that are analogous to word embeddings (Heinzinger *et al*., 2019; Elnaggar *et al*., 2020; Rives *et al*., 2021). These embedding vectors, representing protein sequences, capture the core properties of protein sequences beyond the primary structure, in a way that is context and species agnostic, but relevant to their function in the cell, which makes them particularly useful for understudied organisms Hoarfrost *et al*. (2022). Contextualized word embeddings have already been demonstrated to be successful for predicting GO terms as well as the structure and localization prediction, and refining protein family clusters (Littmann *et al*., 2021; van den Bent *et al*., 2021).

Compared to eukaryotes (Odrzywolek *et al*., 2022), much less has been done to apply NLP-based methods to bacterial function prediction. In a recent CAFA challenge, the competing methods consistently performed worse on bacteria than the eukaryotes, suggesting that there is room for improvement. Furthermore, the prokaryotic track was heavily biased toward a single, well-studied bacterial species, *E. coli* (Zhou *et al*., 2019), pointing to a need to test methodologies on diverse bacteria. Given the vast diversity of functional repertoire in bacterial organisms, remote homology detection is of utmost importance.

Many functionally related bacterial genes are encoded in operons, co-located clusters of genes encoded on the same strand, which are often co-regulated and co-transcribed. Thus, the context of a gene is another means to infer clues to its function (de Daruvar *et al*., 2002; Li *et al*., 2009), as it is a source of functional information which is complementary to both the amino-acid sequence and the embeddings-based representation of a gene. Leveraging gene context and gene interactions as an additional feature was shown to improve prediction performance on eukaryotes (Makrodimitris *et al*., 2020; Yao *et al*., 2021). However, combining the information from gene context with embeddings-based gene representations has not yet been done for function prediction in prokaryotes.

We developed the Synteny-Aware function Predictor (SAP), a novel synteny-aware approach to improve bacterial gene function prediction based on protein embeddings and a comprehensive bacterial operon database. To evaluate SAP, we performed extensive benchmarking using ground truth data and automated function prediction (AFP) standard approaches to show that SAP outperformed conventional sequence-based bacterial genome annotation pipelines, more sophisticated HMM-based approaches, and a state-of-the-art deep learning method when using gene synteny conservation as additional input. As part of a real-world application, we also demonstrated SAP’s utility to predict protein functions in *Enterococcus* species, including predicting potential novel pore-forming toxins related to the delta toxin family that could not be recognized using linear sequence or protein domain information. SAP provides a powerful new tool for protein function prediction in bacteria, combining state-of-the-art NLP methods with a novel incorporation of syntenic information for bacteria.

## 2 Materials and methods

### 2.1 Datasets

#### 2.1.1 SwissProt data set for benchmarking

We retrieved all the manually reviewed entries from the SwissProt Database (release 2021-04, retrieval date 10 November 2021) (The UniProt Consortium, 2018), which was filtered to include proteins of length 40-1000 amino acids and with at least one experimental GO annotation. We selected the evidence codes EXP, IDA, IPI, IMP, IGI, IEP, HTP, HDA, HMP, HGI, HEP, IBA, IBD, IKR, IRD, IC, and TAS. To reduce redundancy, we clustered the proteins using CD-HIT (Li and Godzik, 2006) at 95% sequence similarity. The final dataset comprised 107,818 proteins in total. To benchmark the performance of our method for bacterial gene function prediction, we created five benchmarking datasets from the SwissProt proteins, one for each of the five most numerous bacterial organisms in our final dataset (Table 1). Each organism’s dataset was split into training and test sets. We also divided the full training set in different ways to create five sets where the sequence similarity (calculated using BLASTp (Altschul *et al*., 1990) of test to training set proteins was at most 40%, 50%, 60%, 70% and 80%. This resulted in a total of 30 benchmarking sets (Table 1).

**Table 1.** Total number of proteins in the benchmarking sets generated from the SwissProt dataset to evaluate function prediction tools on bacterial organisms. For each organism, the test set remained constant, whereas the training set was restricted according to the maximum sequence similarity allowed between the test and training sets.

| Organism name | # proteins in the test set | # proteins in the training set<sup>1</sup> |  |  |  |  |  |
| --- | --- | --- | --- | --- | --- | --- | --- |
|  |  | 40% | 50% | 60% | 70% | 80% | Full |
| *Escherichia coli (EC)* | 3454 | 87014 | 96471 | 100445 | 102262 | 103229 | 104377 |
| *Mycobacterium tuberculosis (MT)* | 1666 | 95367 | 102531 | 105158 | 105917 | 106114 | 106152 |
| *Bacillus subtilis (BS)* | 1636 | 93363 | 101112 | 104325 | 105609 | 106015 | 106182 |
| *Pseudomonas aeruginosa (PA)* | 1014 | 94679 | 101338 | 104644 | 106186 | 106680 | 106804 |
| *Salmonella typhimurium (ST)* | 774 | 100928 | 104164 | 105384 | 105980 | 106340 | 107044 |

<sup>1</sup>The maximum sequence similarity between the training and test sets is stated for the first five pairs, and the "Full" column on the far right corresponds to the case where the maximum similarity is 95% to avoid redundancy.

#### 2.1.2 *Enterococcus* diversity dataset

We applied SAP to a set of 61,746 proteins with no experimental annotations, representing the entire protein content of 19 different *Enterococcus* species, spanning four *Enterococcus* clades (Lebreton *et al*., 2017) (Supplementary Table S5). This collection of genomes is representative of *Enterococcus* genomic diversity, hence we refer to it as the *Enterococcus* diversity dataset. Full assemblies were downloaded from the Assembly Database in NCBI (National Library of Medicine (US), 1988).

### 2.2 Building the bacterial operon database, SAPdb

In order to establish a comprehensive, broad compilation of putative conserved bacterial operons in SAPdb to use as a resource for our function prediction tool, we started with the 45,555 representative genomes from the Genome Taxonomy Database (GTDB Release 202, retrieved on 31/03/2022) (Parks *et al*., 2021). We extracted all protein sequences from the standardized GTDB annotations and clustered them using CD-HIT (Li and Godzik, 2006) at 95% sequence identity with default parameters, keeping only the clusters that contained at least 10 genes, resulting in 372,308 clusters of bacterial proteins. Next, we identified operons by grouping together clusters if at least one of the cluster members was located on the same contig and same strand, within 2000 bp (Fig. 1A). This yielded 1,488,249 non-singleton candidate operons. Finally, we removed those with an intergenic distance larger than 300 bp, or split them into multiple operons if possible (Fig. 1B). At the end of this procedure, SAPdb consisted of 406,293 unique non-singleton operons, and the largest operon was 25 genes long.

**Fig. 1.**
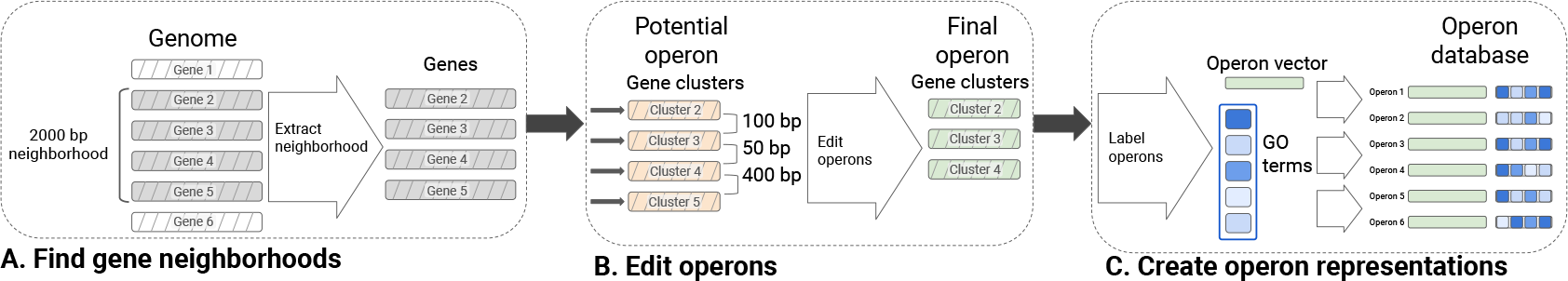
Schematic diagram of method used to construct our operon database, SAPdb. Hashed boxes represent genes; solid boxes are numerical embedding vectors. A. 2000 bp-long gene neighborhoods are extracted from all genomes in GTDB; shown is an example with four genes in a single genomic neighborhood (hashed grey boxes). B. After clustering all proteins from GTDB with CD-HIT, we replace the genes with the CD-HIT clusters they belong to (hashed orange boxes) using the amino-acid sequence of the representative gene of each cluster in place of their actual amino-acid sequence. Then, we trim potential operons to remove genes separated by > 300bp, resulting in final operon clusters (hashed green boxes). C. Once the final operon structures are determined, we i) annotate each operon with a set of GO terms, for which we track the corresponding frequency among the gene clusters that make up the operon (blue rectangles, darker shades mean GO terms are found in more genes within the operon), and ii) extract numerical embedding vectors for each operon (solid green boxes). We create a new representation for each operon, which consists of the average embedding vector and a set of GO terms. The final operon database is a collection of such representative embedding vectors and GO term frequency vectors; representations of six example operons are shown here.

We used experimentally determined operons collected in the Operon DataBase (ODB v4) (Okuda and Yoshizawa, 2010) to help determine threshold values used when building SAPdb, and to validate SAPdb. We downloaded both the ODB known and ODB conserved operon databases on 31/03/2022. We identified operons in ODB belonging to *E. coli* and *B. subtilis*, as (i) these two organisms form the basis of a large part of the benchmarking of SAP, (ii) we could cross-reference the protein IDs in ODB to the locus tags in their respective genome assemblies, and (iii) they are two of the most well-represented organisms in ODB. The ODB conserved operon database contained 8235 unique operons, from which we extracted descriptive statistics and common patterns found across several operons conserved among bacterial organisms. The ODB known operon database was used to model operon features and determine thresholds, such as an operon length, number of genes in an operon, and the maximum intergenic distance between adjacent genes in an operon.

To summarize each SAPdb operon, we extracted the protein embedding vectors for the representative protein sequence of clusters found in that operon. We used ESM-1b, a transformer-based protein language model (Rives *et al*., 2021) to extract the embeddings, and we took the average of these embeddings to obtain one embeddings vector per operon (Fig. 1C). Then, we annotated the operons in SAPdb by assigning GO terms, if possible. Since we did not have experimental annotations, we labeled operons based on sequence similarity. We used BLASTp (Altschul *et al*., 1990) to calculate pairwise sequence similarity between proteins in SAPdb operons and the non-redundant SwissProt database with experimentally determined GO terms (all 107,818 entries). We transferred GO terms found in significant hits (e-value < 1e-6 and bit score > 50) using the frequency of each GO term among these hits as a predicted score. With this approach, we could assign at least one GO term to 295,446 of the 372,308 clusters of bacterial proteins (79%), which in turn yielded 388,377 non-singleton operons (out of 406,293; 96%) annotated with at least one GO term (Table S2).

In order to keep our operon database consistent with our benchmarking datasets, where we evaluated SAP on training subsets with differing sequence similarity to the proteins in the test set, we generated corresponding subsets of SAPdb with matching sequence similarity thresholds. We followed the same procedure as we did to generate subsets of the SwissProt training sets with different sequence similarity thresholds: we used BLAST to calculate the pairwise sequence identity of each query protein to the protein clusters that form our main operon database. We removed clusters if they were more than 40%, 50%, 60%, 70%, 80% and 95% similar to at least one of the query proteins in the test set. Since this operation removed or altered the content of operons, we re-calculated the intergenic distances for the remaining clusters and again split operons where the intergenic distance exceeded our 300bp threshold, as we did when we created the main operon database (Fig. 1B-C).

### 2.3 Comparison to published function prediction methods

#### 2.3.1 Comparison to broadly used function prediction methods as baseline

In our SwissProt benchmarks, we compared SAP to two conventional function predictors: i) BLAST (v. 2.12.0) (Altschul *et al*., 1990), a standard sequence homology-based predictor used widely in the literature for comparisons, and ii) an HMM-based approach, which serves as a more sophisticated baseline.

To predict function using the BLAST baseline, we transferred GO terms from significant BLAST hits (e-value < 1e-3) of a query protein with a predicted score of the value of the maximum sequence identity.

As an alternative, we also used the GO term frequency-based approach (Zhou *et al*., 2019; van den Bent *et al*., 2021), but we found the maximum sequence identity scoring method performed better in our experiments.

To predict function using the HMM-based approach, we ran HMMER (Eddy, 2011) against the Pfam database and applied the frequency-based approach to score transferred annotations, i.e. we transferred GO terms from all significant HMM hits (e-value < 1e-3) to the query protein, using the frequency of a GO term (number of instances a term was observed among the significant hits) as the predicted score. To compare Pfam outputs quantitatively with the rest of the methods, we used the mapping tables provided by the GO consortium to obtain GO terms corresponding to each Pfam ID (Ashburner *et al*., 2000). Since the Pfam database is independent of the train/test pairs we generated for our experiments, we report the same numerical results for all pairs.

#### 2.3.2 Comparison to current state-of-the-art tools based on deep learning methods

We also compared SAP to two newer tools based on deep learning methods. We first opted for a simple, unsupervised approach (which we will call the knn approach). We used the ProtT5-XL-U50 (which we will call T5) (Elnaggar *et al*., 2020) and ESM-1b (which we will call ESM) (Rives *et al*., 2021) protein language models to represent protein sequences. To extract amino-acid level embedding vectors, we used `bio_embeddings` (v 0.2.2) (Dallago *et al*., 2021) with default settings. Then, we obtained protein-level embeddings (1024 dimensional vectors for T5 and 1280 for ESM) by taking the average over individual amino acid embeddings to obtain embedding representation vectors. In preliminary experiments and our current benchmark study, we found that the embeddings extracted from the ESM model performed better; thus we use only the ESM model throughout this work.

In order to transfer annotations using these embeddings, we used a nearest neighbor predictor (named knn), which was designed in a similar manner to goPredSim (Littmann *et al*., 2021). For each query protein, we identify nearest neighbors in the training set based on embedding vector similarity over a given threshold, which we calculate separately for each query as the x^th^ percentile among all pairwise similarity values, where x parameter is set to 99 percentile. We transfer GO terms from the nearest neighbors with a score equal to their cosine similarity to the query protein. As the final prediction, we keep only the maximum score for each GO term transferred from the nearest neighbors. Throughout this work, we use cosine similarity to determine the similarity between any two embedding vectors 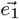 and 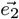.

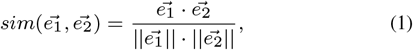

where 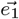 and 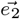 are both real-valued vectors, 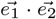 represents the dot product between 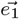 and 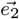, and 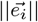 is the Euclidean norm of vector 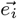, where i = 1, 2.

We chose DeepGOPlus (v 1.0.1) (Kulmanov and Hoehndorf, 2019) as the second deep learning based competitor in our experiments. DeepGOPlus, one of the state-of-the-art tools in the field, is a supervised approach where a deep convolutional neural network model is combined with a sequence homology based method. We used the DeepGOPlus implementation provided by the authors; we trained the model on the training sets in our experiments with the optimal values reported for the hyperparameters (Kulmanov and Hoehndorf, 2019). We used the same training set for both the BLAST queries and the deep learning based methods.

### 2.4 SAP: Synteny-aware function prediction using protein embeddings

SAP combines protein embeddings to represent amino-acid sequences by leveraging conserved synteny among bacterial operons to help identify function prediction, in two main steps (Fig. 2): (i) assigning operons to a query from the pre-computed bacterial operon database, SAPdb (Fig. 2A) and (ii) transferring GO terms from SAPdb operons to the query (Fig. 2B).

**Fig. 2.**
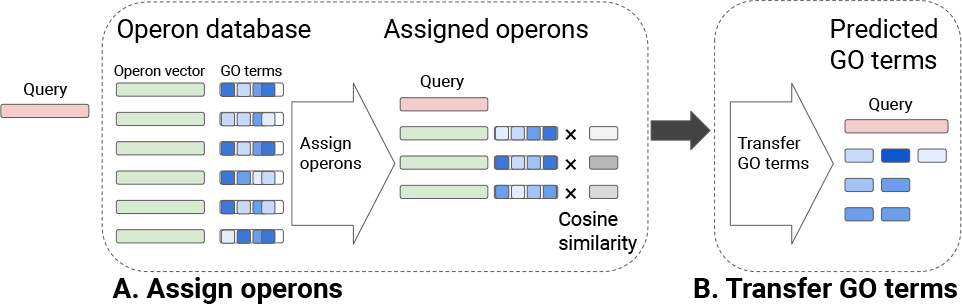
Overview of SAP algorithm: predicting GO terms of a query protein. A. SAP assigns an operon (or multiple operons) to the query protein (red filled rectangle on the left) represented using embeddings from ESM-1b LM, based on cosine similarity. Consistent with Fig. 1, green rectangles show operon embeddings paired with the corresponding GO term frequencies (blue rectangles). In this example, three operons that passed the threshold are assigned to the query, and their GO term frequencies are weighted by multiplying by the cosine similarity. B. All GO terms from the assigned operons are transferred to the query, where the final predicted score of a GO term is the maximum of all the multiplied values for the term.

For each query, we identify the most suitable operons in our database following the same procedure as we did for the nearest neighbor predictors based on protein embeddings, and we use the ESM model to extract the embedding vectors. In short, we calculate the pairwise cosine similarity between the query point and the average embedding vectors representing operons in the database. We assign an operon to the query if the pairwise similarity between the operon embeddings and the query embeddings is greater than a threshold, corresponding to the x^th^ percentile among all pairwise similarity values. In our current implementation, we do not have any restrictions on operons assigned to a query protein: given that the most suitable operons are picked among the same set of operons used to calculate the threshold, at least one operon is assigned to each query point. For all such operons assigned to the query, we also retrieve the GO term frequencies. We transfer all the GO terms found in the assigned operons using the frequency of the terms multiplied by the cosine similarity of the query point to the operon as the predicted score. For each GO term, the predicted score is the maximum of these values. As the final step in our algorithm, we normalize the predicted scores separately within three GO classes. In this work, SAP uses our operon database, SAPdb (Fig. 1).

In addition to using SAP as described above, we also tested running a version of SAP which uses only the operon database (titled SAP-operon). Evaluating these two methods side by side allows us to assess the contribution of using our operon database on SAP’s performance. For SAP-operon, we removed all singleton entries from the database and relied only on those that were at least two genes long; thus, all gene predictions will originate from the operons.

### 2.5 SwissProt benchmark evaluation

Using our SwissProt benchmarking datasets, we evaluate six different protein prediction methods: two baselines (BLAST and Pfam), a nearest neighbor predictor based on protein embeddings extracted using the ESM-1b model (knn), SAP and its variant SAP-operon, where only the operon component of SAP is retained, and DeepGOPlus. In order to make the outputs of all tools comparable to those of DeepGOPlus, we propagated the predicted GO term scores based on the GO hierarchy, following the procedure in (Kulmanov and Hoehndorf, 2019). For each GO term, we assigned the highest predicted score from among all its children. This additional post-processing step was only implemented in our benchmarking comparisons across tools, and not in our function prediction across the *Enterococcus* genus.

We evaluated these function prediction methods as done for the CAFA challenges, using the maximum F1-score (*F*_max_) and the minimum semantic distance (*S*_min_) as described in (Radivojac *et al*., 2013). We also report the coverage, defined as the percentage of test proteins annotated with at least one GO term at the threshold which maximizes the F1-score.

### 2.6 Applying SAP to a diverse set of enterococcal genomes, including detailed analysis of pore-forming toxins

To demonstrate a practical application of SAP, we applied SAP to the *Enterococcus* diversity dataset. We ran SAP in default mode on this full set of genes, and compared the output to that from three other bacterial gene annotation approaches: (i) the prokka annotation pipeline (v. 1.14.6) (Seemann, 2014), which runs multiple sequence homology-based function prediction tools; (ii) the Pfam database (release 32.0) (Paysan-Lafosse *et al*., 2023) using HMMER (v 3.3.2) (Eddy, 2011); and (iii) eggNOG mapper (v 2.1.10) (Huerta-Cepas *et al*., 2018; Cantalapiedra *et al*., 2021). All tools were run using default parameter settings; for both HMMER and eggNOG mapper, a significant hit was defined as having e-value < 1e-3.

When examining potential novel *Enterococcus* pore-forming toxins, we performed additional analyses to assess the potential function of query proteins without experimental annotations: (i) we performed a large-scale structure search using the query protein against AlphaFoldDB and the Protein Data Bank (PDB); (ii) we examined their similarity to known pore-forming toxins found in *Enterococcus* or closely related genera (Table 2), both in terms of structural similarity (using Foldseek), as well as in genomic context; and (iii) we assessed the presence of key structural elements, including N-terminal signal sequences, a common feature in most toxin sequences which guides toxin secretion and transportation outside the cell. In order to compare syntenic relationships between predicted and known toxin genes, we examined five genes upstream and downstream of toxin genes predicted by SAP, as well as for the known delta toxin genes from Table 2, *epx1* and *epx4* (Xiong *et al*., 2022).

**Table 2.**
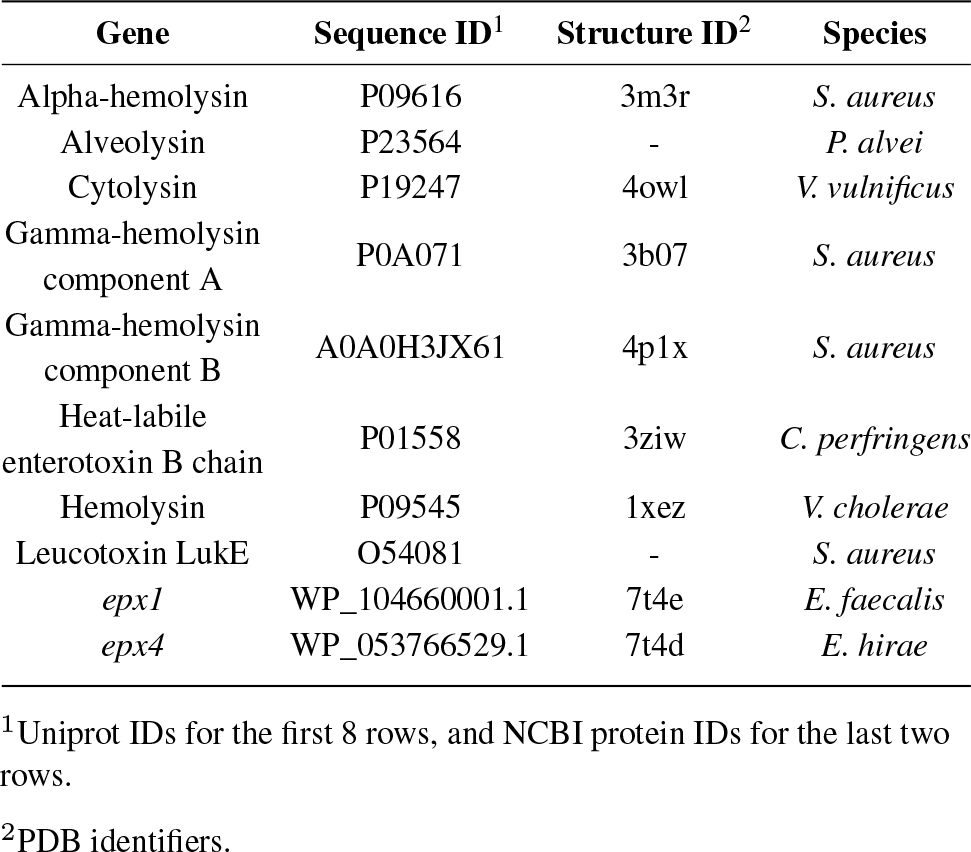
Known pore-forming toxin genes found in Enterococcus or closely related genera. Protein structures for all but two genes have been solved experimentally and were downloaded from PDB. For Leucotoxin and Alveolysin, AlphaFold was used to predict their structure since they were not found in the databases.

To predict the structure of potential novel toxin genes identified by SAP, we used the Fold Sequence public server on ESMFold Atlas (Lin *et al*., 2023) which only allows input sequences shorter than 400 amino acids. For longer proteins, we used AlphaFold (Jumper *et al*., 2021). We ran AlphaFold in monomer mode with default settings using the Docker implementation. We used Foldseek (van Kempen *et al*., 2023) for both protein structure search against databases and structural alignment. While the structure database search was performed with default settings, we utilized both the global (`--alignment-type 1`) and local alignment options (`--alignment-type 2`) of Foldseek. Following the guidelines available for running Foldseek, we labeled alignments as highly significant (structural alignment score > 0.7), significant (0.6 < structural alignment score *≤* 0.7), nonrandom (0.5 < structural alignment score *≤* 0.6) or random (structural alignment score *≤* 0.5). To account for large differences in the query and target sequence length, we required the alignment probability to be greater than 0.8 as well. We predicted the N-terminal signal sequences using the SMART server (Schultz *et al*., 1998).

## 3 Results

To improve gene function annotation for bacteria, we developed SAP, which combines state-of-the-art protein embeddings based on NLP algorithms with bacteria-specific information about the gene function inferred from conserved bacterial operons. In brief, SAP represents amino acid sequences using NLP-based embedding vectors and calculates protein similarity using the distances between embedding vectors. Our custom-built database, SAPdb, an extensive collection of bacterial operons and their annotations, is used to find operons containing proteins similar to the query. Annotations from these operons are transferred to the query, allowing us to leverage additional functional information from similar genomic neighborhoods in other organisms. SAP uses the ESM-1b protein language model, together with a k-nearest neighbors (knn) framework for transferring GO terms (Methods), which we found to be the best option because it achieved the highest prediction scores in our benchmarks.

### 3.1 SAPdb: an operon database to leverage functional information derived from syntenic relationships across bacteria

To incorporate information about operon structure into SAP, we constructed a large-scale database, which we named SAPdb, of over 400,000 operons predicted from > 45,000 representative genomes from across the bacterial kingdom (Methods). We validated SAPdb by comparison to the experimentally determined operons found in the conserved Operon DataBase (ODB) (Okuda and Yoshizawa, 2010), a similar online database. SAPdb is larger-scale and more up-to-date than ODB, which is based on a smaller, curated list of experimentally determined operons from the literature. Overall, SAPdb, is quantitatively similar to the conserved ODB, in terms of operon length, number of genes in an operon and intergenic distance within operons (Supplementary Fig. S1-S3). SAPdb provides an extensive catalogue of conserved patterns of gene synteny within the bacterial kingdom (Table S1).

### 3.2 SAP outperforms other tools in function prediction for multiple bacterial species

To assess the performance of SAP in assigning GO terms to proteins, we first performed benchmarking on the SwissProt database, where only the proteins with at least one experimentally determined GO annotation were retained. We then created benchmarking datasets for five different bacterial species, dividing SwissProt entries into training and test sets, thus simulating the real-world scenario of annotating predicted proteins that lack exact matches to database entries.

We benchmarked SAP against four tools, including i) a baseline BLAST method; ii) a basic HMM-based approach (HMMER); iii) a simple, unsupervised deep learning method (knn); and iv) a state-of-the-art deep learning method (DeepGOPlus) (Methods). We performed benchmarking separately for three categories of GO terms, including Biological Process (BPO), Molecular Function (MFO), and Cellular Component (CCO), as these three categories are known to present different challenges for annotation (Radivojac *et al*., 2013). Overall, SAP achieved the highest F_max_ scores across all five bacterial species, for all three GO categories, and on the full SwissProt benchmarking set, with *S. typhimurium* being the only exception. On this species, DeepGOPlus performed the best for BPO and MFO (Table 3). We observed similar trends in prediction performance using S_min_ and the area under the precision/recall curve (Supplementary Tables S5 and S6).

**Table 3.** *F* _max_ scores from our benchmarking for six different function prediction tools, for each of five bacterial species in our full SwissProt benchmarking set. *F* _max_ scores are tabulated separately for the three GO categories, BPO, MFO and CCO. The highest *F* _max_ score in each column is shown in bold.

| Method | Bacterial species<sup>1</sup> |  |  |  |  |
| --- | --- | --- | --- | --- | --- |
|  | EC | MT | BS | PA | ST |
| $F_{\max}$ scores for BPO | | | | | |
| BLAST | 0.570 | 0.543 | 0.639 | 0.683 | 0.852 |
| Pfam | 0.610 | 0.513 | 0.582 | 0.579 | 0.579 |
| knn | 0.646 | 0.636 | 0.828 | 0.797 | 0.880 |
| DeepGOPlus | 0.648 | 0.669 | 0.857 | 0.824 | **0.928** |
| SAP-operon | 0.872 | 0.837 | **0.915** | 0.928 | 0.903 |
| SAP | **0.876** | **0.838** | **0.915** | **0.929** | 0.902 |
| $F_{\max}$ scores for MFO | | | | | |
| BLAST | 0.613 | 0.593 | 0.625 | 0.699 | 0.814 |
| Pfam | 0.650 | 0.549 | 0.571 | 0.534 | 0.559 |
| knn | 0.675 | 0.723 | 0.814 | 0.854 | 0.837 |
| DeepGOPlus | 0.686 | 0.755 | 0.841 | 0.883 | **0.911** |
| SAP-operon | 0.880 | **0.869** | **0.893** | **0.938** | 0.878 |
| SAP | **0.885** | **0.869** | **0.893** | **0.938** | 0.877 |
| $F_{\max}$ scores for CCO | | | | | |
| BLAST | 0.569 | 0.397 | 0.638 | 0.700 | 0.871 |
| Pfam | 0.625 | 0.541 | 0.608 | 0.560 | 0.616 |
| knn | 0.731 | 0.500 | 0.898 | 0.900 | 0.917 |
| DeepGOPlus | 0.745 | 0.567 | 0.885 | 0.887 | **0.936** |
| SAP-operon | 0.920 | **0.847** | **0.943** | **0.945** | 0.918 |
| SAP | **0.922** | **0.847** | **0.943** | **0.945** | 0.917 |

<sup>1</sup>EC: *Escherichia coli*, MT: *Mycobacterium tuberculosis*, BS: *Bacillus subtilis*, PA: *Pseudomonas aeruginosa* and ST: *Salmonella typhimurium*.

Also, when protein embeddings were used, even in a simple unsupervised setting (such as knn), they provided a better representation of protein sequence for GO term transfer than both the amino-acid sequence itself (BLAST baseline) and the HMM profiles (Pfam baseline) (Table 3). This agreed with recent studies on eukaryotes (Heinzinger *et al*., 2022).

### 3.3 SAP surpasses existing tools for detection of distant homologs

We were particularly motivated to develop SAP to increase the number of annotations for the very large number of novel proteins of completely unknown function within bacteria, which occurs when a predicted protein has no or very low homology to existing databases. To emulate gene function prediction in such low homology instances, we designed additional benchmarking sets where the pairs of training and test sets were generated by stratifying the full SwissProt dataset based on the maximum sequence similarity allowed between protein sequences in the training and the test set.

For each of the five bacterial species, we constructed additional benchmarking sets with five sequence similarity thresholds: 40%, 50%, 60%, 70%, and 80%, where 40% or less sequence similarity presents the most challenging scenario for annotation. For consistency and to minimize the chance that the operons could artificially inflate SAP’s performance, i.e. information leak between the training and test sets, we also modified our operon database to remove clusters homologous to the test sequences and, for each pair of training and test sets, we rebuilt the operon database consistent with its homology threshold. NLP-based embeddings (knn) far outperformed both conventional predictors, BLAST and Pfam, across the whole range of sequence similarities. As we did not observe any significant differences between the species examined, we report the average F_max_ values and standard deviation for all five bacteria combined (BPO in Fig. 3). SAP was consistently the top-performing method. The difference in prediction performance (as measured by F_max_) between SAP and all other methods was greater as the sequence similarity between the test and the training sequences (as well as the clusters in the operon database) increased (Fig 3).

**Fig. 3.**
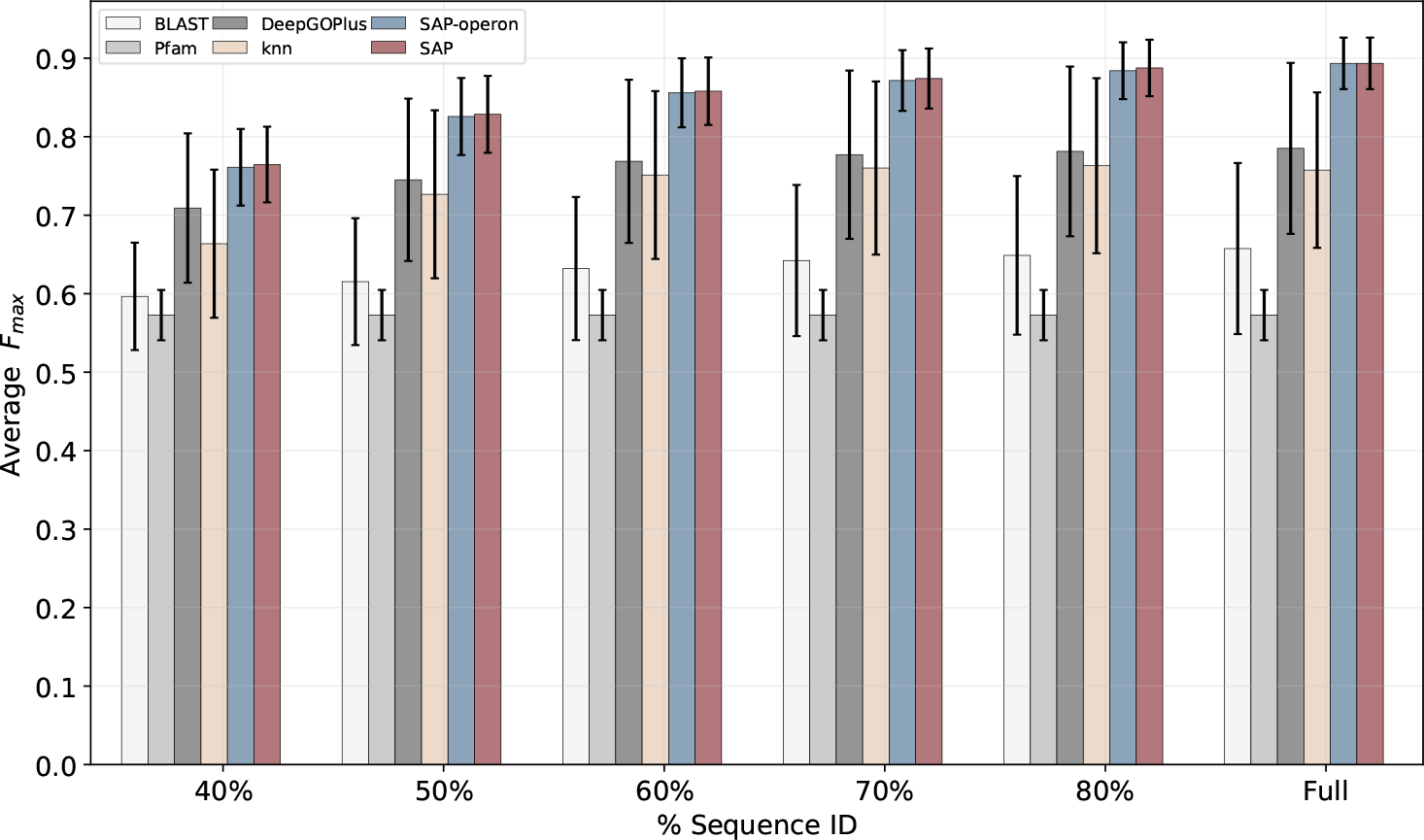
SAP outperformed conventional approaches to function prediction, averaged across five different bacterial species, for varying levels of protein sequence identity to the proteins in the training set (x-axis). The error bars show the standard deviation for each method. The *F*_max_ values for the Pfam baseline are identical for all 6 experiment sets. Because Pfam uses a different training set, we were unable to modify the % sequence identity input. Thus, for Pfam we only plot a single value per species.

In addition, this benchmarking revealed that BLAST performance was surprisingly consistent over different levels of shared homology while the embeddings-based methods all showed incremental improvement in performance as homology between the training and test sets increased. This trend held for not only the average F_max_ in the remaining two ontologies (MFO and CCO), but also for each bacterial species individually (Supplementary Tables S8-S12).

### 3.4 SAP provides more reliable predictions compared to other methods

Among the tools in our benchmark, the BLAST and HMM-based Pfam baselines had the lowest annotation coverage values (i.e. the number of test genes that have at least one predicted GO term) on both the full SwissProt dataset and the remote homology detection tests (Tables S13 and S14). SAP emerged as the all-around top-performing method in terms of balancing precision and recall. Furthermore, we found that its prediction coverage was in line with other embeddings-based knn models on the full SwissProt benchmarking, although it occasionally lagged behind the state-of-the-art in terms of coverage on our other benchmark sets. Given that SAP achieved the best *F*_max_ values across the board, the drop in coverage means SAP’s predictions are more reliable compared to other methods in our benchmark. We did observe that SAP’s coverage decreased slightly for test sets with lower homology to the training set (Table S14). In these low homology tests, SAPdb is sparsely labeled due to a conservative annotation methodology (Supplementary Text), limiting the annotations that can be transferred based on synteny.

### 3.5 Applying SAP to a diverse set of enterococcal genomes, including identification of five potential novel pore-forming toxins

A key goal in the development of SAP was annotating novel genes of unknown function, including those associated with key bacterial features of clinical interest such as antimicrobial resistance and virulence. *Enterococcus* is a diverse genus of bacteria thought to inhabit the gastrointestinal tracts of all land animals. These organisms have an incredibly diverse functional repertoire, yet many of their predicted proteins are of unknown function (Lebreton *et al*., 2017; Schwartzman *et al*., 2023). Uncovering this rich functional diversity is of primary interest given the ubiquity and importance of this genus. Recent targeted searches have reported the discovery of several classes of novel toxins within diverse enterococcal species, including the discovery of a new family of pore-forming delta toxins in *E. faecalis, E. faecium* and *E. hirae* (Xiong *et al*., 2022) and new botulinum toxins in *E. faecium* (Zhang *et al*., 2018). All of these newly discovered toxins exhibit low sequence homology to known toxin sequences in other bacterial species.

Although the previous studies focused only on three clinically relevant species of *Enterococcus*, we hypothesize that similar toxins could also be found in other diverse, less well-studied species of *Enterococcus*, providing insights into other ecologies in which these toxins may be advantageous. Thus, to search for additional novel toxin genes across the *Enterococcus* genus, we applied SAP to a collection of 19 *Enterococcus* genomes, each representing a different species (Lebreton et al., 2017), including 16 species not examined by Xiong *et al*. or Zhang *et al*.. We looked specifically for genes that were labeled with a GO term describing toxin activity and associated with the conserved genomic context of delta toxins (Xiong *et al*., 2022). SAP associated 59 genes with the single delta toxin operon from SAPdb, consisting of an enterotoxin and a putative lipoprotein cluster, found in the unrelated *Clostridium* and *Roseburia* species (Table S4). Of these 59 genes, 6 were predicted by Pfam to be pore-forming toxins (e-value < 1e-3 to PF01117 or PF03318), and 3 were annotated by Prokka as “lipoproteins” (Methods). The remaining 50 had no functional prediction prior to running SAP.

To explore their candidacy as delta toxin encoding, we evaluated each gene’s predicted protein structure and genomic context. Eleven (of 59) had structural similarity to known toxin structural folds (Foldseek alignment probability > 0.8 and alignment score > 0.5 to proteins in the AlphaFold and the Protein Data Bank (PDB) structure databases), including several with highly significant alignments (Table 2; Fig. 4). Of these eleven, five were not previously identified as having a toxin annotation by either Prokka or Pfam - these were detected only by SAP. All 11 of these genes contained signal peptides at similar positions as those in known bacterial toxins. The remaining 48 proteins without structural similarity had lower SAPdb rankings than the 11 with structural similarity (Supplementary Text).

**Fig. 4.**
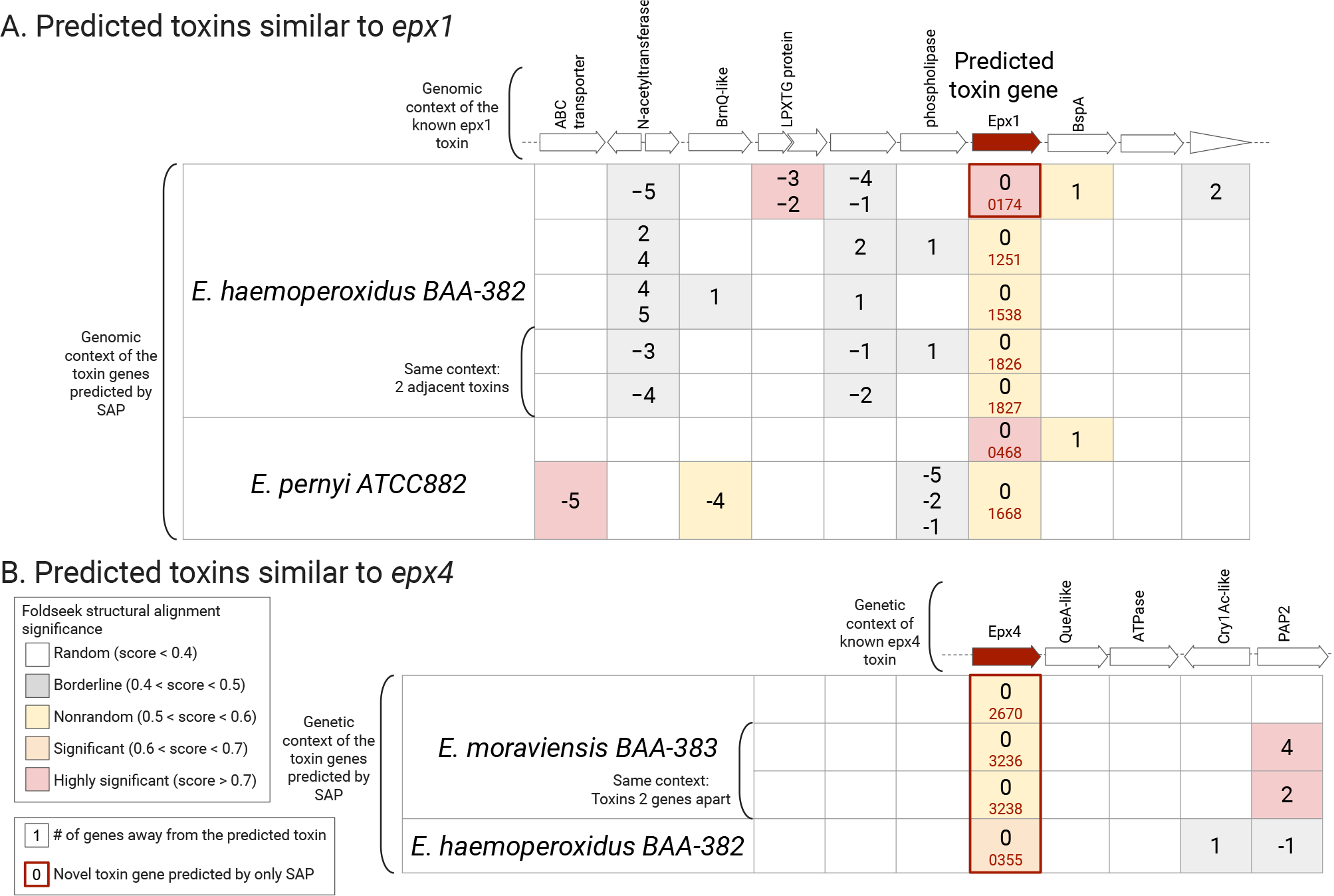
SAP predicts 11 likely novel toxin genes in our Enterococcus dataset (aligned below the column titled “Predicted toxin gene”), including five that could not be predicted by previous function prediction methods (red frames). A) seven genes with the highest similarity to E. faecalis epx1. These genes also shared some similarity in genomic context with epx1. B) four genes with the highest similarity to E. hirae epx4. These genes also shared some similarity in genomic context with epx4. The operon diagrams show the known genomic contexts of epx1 and exp4 found in E. faecalis and E. hirae species that were studied, respectively (Xiong et al., 2022). Beneath this, cells in the table represent the occurrence of genes with structural similarity to those in the known epx1 or epx4 operons. Their relative position within the operon in reference to the predicted toxin gene (# of genes away from the predicted toxin) and structural alignment score (coloring) obtained using Foldseek to the analogous gene in the operon diagram is shown. Gene locus tags, a 4-digit number given based on their location within the genome, for the predicted toxins are also placed within the cells. Additional details are shown in Figure S7.

We compared the genomic context of the 11 candidate toxins identified by SAP to the known pore-forming delta toxin genes previously reported in *Enterococcus, epx1* and *epx4*, and their neighborhoods (Xiong *et al*., 2022). Seven of the 11 candidate toxin genes were most similar to *epx1* structures from *E. faecalis* and *S. aureus*, including five from *E. haemoperoxidus* BAA-382, and two from *E. pernyi* ATCC882. All had surrounding genes with some degree of structural similarity to genes within the known *epx1* genomic neighborhood, including two with highly significant matches (Fig. 4A). Among the 5 putative toxin genes, the highest amino acid sequence identity to *epx1* was less than 40% (Supplementary Table S15). Furthermore, the gene neighborhood was conserved between the five candidates from *E. haemoperoxidus* BAA-382 (Fig. 4).

Four of the 11 candidate toxin genes were most similar to the *E. hirae epx4* structure, including one gene from *E. haemoperoxidus* and three genes from *E. moraviensis* BAA-383 (Fig. 4B). Among the four putative *epx4* genes, the maximum amino acid sequence similarity we observed to *epx4* was 60% (Supplementary Table S15). Similar to the *epx1* context, we observed that the neighboring genes of the new *epx4*-like toxins predicted by SAP were structurally similar to one another. Although some of the neighboring genes had lower Foldseek similarity scores, the neighborhoods had nonrandom similarity among themselves (scores ranging from 0.4 to 0.9). Thus, SAP detected novel toxins, found in a conserved genomic context, that other tools could not identify. The combination of structural similarity with commonalities in the genomic neighborhood makes these genes interesting targets for further experimental validation.

## 4 Discussion

In this work, we introduce SAP, a novel synteny-aware, NLP-based function prediction tool for bacteria. SAP is distinguished from existing tools for annotating bacteria in two ways: (i) it represents proteins using embedding vectors extracted from state-of-the-art protein language models, and (ii) it incorporates additional functional information inferred from a protein’s genomic neighborhood, by leveraging conserved synteny across the entire bacterial kingdom, tabulated in our operon database SAPdb. To our knowledge, SAP is the only bacterial gene function prediction tool with these two features.

While there have been several successful uses of protein language models for protein function prediction in eukaryotes, these methods have not yet been extensively applied to bacterial organisms (Littmann *et al*., 2021; Yao *et al*., 2021). We used embedding vectors in SAP, motivated by recent work showing significant improvements in gene function prediction by replacing features derived from amino-acid sequences with embedding vectors extracted from protein language models. To assess SAP’s performance on bacteria, we designed a systematic, rigorous experimental framework based on the SwissProt database where we evaluated function predictors for remote homology detection as well. We confirmed that protein embeddings surpass conventional sequence homology-based tools across diverse bacteria, and that they provide a better representation of genes to infer gene function (Table 3).

Although bacterial gene neighborhoods have been used previously for function prediction, this practice has mostly been manual and is absent from current automated annotation tools. We designed a purely computational, bottom-up approach to incorporate this syntenic information into bacterial gene function prediction, thereby leveraging this valuable source of functional information. We demonstrate that conserved synteny and protein embeddings provide complementary information for predicting gene function, in particular for remote homology detection, as demonstrated via our systematic, rigorous experimental framework to evaluate function predictors based on the SwissProt database (Fig. 3). We consistently achieved the best performance when operons were used in conjunction with the embeddings representation within the SAP framework. Either component alone resulted in lower performance.

Finally, to explore the performance of SAP on a more practical, real-life application, we used SAP to annotate a set of 19 *Enterococcus* species, representing the phylogenetic range of this genus. Following the recent discovery of several different types of novel toxin genes in enterococci (Zhang *et al*., 2018; Xiong *et al*., 2022), we focused on toxin discovery. SAP predicted 11 candidate delta toxin genes, which showed low sequence homology to known toxins (< 30%) but showed significant structural homology to known toxin protein structural folds. Several of these candidates also shared similar genomic neighborhood patterns with those of known toxin genes. Although six of these candidate toxins could also be identified based on their Pfam domains, five of these could not be annotated using any of the existing gene prediction tools. We assert that these genes are strong candidates for further experimental validation of their toxin activities.

One limitation of SAP is its reliance on a predicted operon database, which may contain syntenic linkages which do not share a function. In the absence of ground truth, both the operon predictions and the functions we assigned to these operons are limited by the existing databases (Supplementary Text). To minimize false positives in operon annotations, we adopted a conservative approach which in turn resulted in a sparsely annotated training set, lowering the prediction coverage of SAP (Supplementary Tables S12 and S13). One way to alleviate this problem would be to routinely pick unlabeled operons from our database, prioritizing the most common ones, to perform experiments and identify their functions. With each new experimental annotation available, additional operons can be labeled. We expect this iterative approach to rapidly increase the number of labeled operons available in the database. Another limitation of the current version of SAPdb is its focus on broadly conserved patterns; it represents conserved synteny across the entire bacterial kingdom. Since our goal was to develop an all-purpose bacterial gene annotation tool, we deliberately designed our operon database to be inclusive and to cover as many conserved syntenic regions as possible. Thus, patterns or operons associated with rare traits in bacteria, or functional pathways unique to novel species are not present in the default SAPdb, but are straightforward to add for specific analyses. Currently, SAP assigns every query gene the same number of operons, equal to 1% of all operons available in the dataset, which we opted to be as inclusive as possible in learning about previously unannotated genes. To help disambiguate real matches from false positive matches, SAP reports a rank for each of the matching operons based on their similarity to the query gene. While we have not determined whether a universal ranking threshold exists, our detailed examination of toxin operons in *Enterococcus* suggested this ranking can be a reliable proxy for confidence. While SAP reported 48 additional genes associated with the delta toxin operon, the delta toxin operon only ranked among the top two operons for only the 11 candidate genes that showed structural similarity to the toxin fold. Thus, the order of assigned operons could potentially be used as a proxy to infer confidence in these assignments.

We demonstrated that conserved synteny and protein embeddings both provide useful information for predicting the protein function; SAP outperforms conventional sequence-based bacterial genome annotation pipelines, as well as more sophisticated HMM-based approaches and more recently developed deep learning methods. SAP can not only infer beyond the linear sequence, at the level of protein fold, but it can also successfully utilize conserved synteny among bacterial species to predict gene function.

## Acknowledgements

We thank Stavros Makrodimitris for providing valuable insight and engaging in fruitful discussions at the inception of this work; Tamim Abdelaal for giving compelling feedback. We also thank our collaborators: Gilmore Lab at Harvard Medical School, and the Bacterial Genomics Group at Broad Institute of MIT and Harvard for helpful discussions of the evolution and biology of enterococci.

## Data Availability

All data used for the analyses in this article are publicly available in the Uniprot database and the Assembly Database at NCBI (accession IDs for the *Enterococcus* assemblies can be found in Supplementary Table S5). The code and the scripts developed in this work are public at https://github.com/AbeelLab/sap.

## References

Altschul, S. F. et al. (1990). Basic local alignment search tool. Journal of molecular biology, 215(3), 403–410.

Ashburner, M. et al. (2000). Gene ontology: tool for the unification of biology. Nature genetics, 25(1), 25–29.

Buchfink, B. et al. (2021). Sensitive protein alignments at tree-of-life scale using diamond. Nature methods, 18(4), 366–368.

Cantalapiedra, C. P. et al. (2021). eggNOG-mapper v2: Functional Annotation, Orthology Assignments, and Domain Prediction at the Metagenomic Scale. Molecular Biology and Evolution, 38(12), 5825–5829.

Dallago, C. et al. (2021). Learned embeddings from deep learning to visualize and predict protein sets. Current Protocols, 1(5), e113.

de Daruvar, A. et al. (2002). Analysis of the cellular functions of escherichia coli operons and their conservation in bacillus subtilis. Journal of molecular evolution, 55, 211–221.

Eddy, S. R. (2011). Accelerated profile hmm searches. PLoS computational biology, 7(10), e1002195.

Elnaggar, A. et al. (2020). ProtTrans: Towards Cracking the Language of Life’s Code Through Self-Supervised Deep Learning and High Performance Computing.

Heinzinger, M. et al. (2019). Modeling aspects of the language of life through transfer-learning protein sequences. BMC Bioinformatics, 20(1), 723.

Heinzinger, M. et al. (2022). Contrastive learning on protein embeddings enlightens midnight zone. NAR Genomics and Bioinformatics, 4(2). qac043.

Hoarfrost, A. et al. (2022). Deep learning of a bacterial and archaeal universal language of life enables transfer learning and illuminates microbial dark matter. Nature communications, 13(1), 2606.

Huerta-Cepas, J. et al. (2018). eggNOG 5.0: a hierarchical, functionally and phylogenetically annotated orthology resource based on 5090 organisms and 2502 viruses. Nucleic Acids Research, 47(D1), D309–D314.

Jensen, L. J. et al. (2003). Prediction of human protein function according to Gene Ontology categories. Bioinformatics, 19(5), 635–642.

Jumper, J. et al. (2021). Highly accurate protein structure prediction with AlphaFold. Nature, 596(7873), 583–589.

Kulmanov, M. and Hoehndorf, R. (2019). DeepGOPlus: improved protein function prediction from sequence. Bioinformatics, 36(2), 422–429.

Lebreton, F. et al. (2017). Tracing the enterococci from paleozoic origins to the hospital. Cell, 169(5), 849–861.

Li, W. and Godzik, A. (2006). Cd-hit: a fast program for clustering and comparing large sets of protein or nucleotide sequences. Bioinformatics, 22(13), 1658–1659.

Li, X. et al. (2009). Gene function prediction with gene interaction networks: a context graph kernel approach. IEEE Transactions on Information Technology in Biomedicine, 14(1), 119–128.

Lin, Z. et al. (2023). Evolutionary-scale prediction of atomic-level protein structure with a language model. Science, 379(6637), 1123–1130.

Littmann, M. et al. (2021). Embeddings from deep learning transfer go annotations beyond homology. Scientific reports, 11(1), 1–14.

Makrodimitris, S. et al. (2020). A thorough analysis of the contribution of experimental, derived and sequence-based predicted protein-protein interactions for functional annotation of proteins. Plos one, 15(11), e0242723.

National Library of Medicine (US) (1988). National Center for Biotechnology Information (NCBI). Bethesda (MD). Available from: https://www.ncbi.nlm.nih.gov/.

Odrzywolek, K. et al. (2022). Deep embeddings to comprehend and visualize microbiome protein space. Scientific Reports, 12(1), 1–15.

Okuda, S. and Yoshizawa, A. C. (2010). Odb: a database for operon organizations, 2011 update. Nucleic acids research, 39(suppl_1), D552–D555.

Parks, D. H. et al. (2021). Gtdb: an ongoing census of bacterial and archaeal diversity through a phylogenetically consistent, rank normalized and complete genome-based taxonomy. Nucleic Acids Research, 50(D1), D785–D794.

Paysan-Lafosse, T. et al. (2023). Interpro in 2022. Nucleic Acids Research, 51(D1), D418–D427.

Radivojac, P. et al. (2013). A large-scale evaluation of computational protein function prediction. Nature Methods, 10(3), 221–227.

Rives, A. et al. (2021). Biological structure and function emerge from scaling unsupervised learning to 250 million protein sequences. Proceedings of the National Academy of Sciences, 118(15), e2016239118.

Schultz, J. et al. (1998). Smart, a simple modular architecture research tool: Identification of signaling domains. Proceedings of the National Academy of Sciences, 95(11), 5857–5864.

Schwartzman, J. A. et al. (2023). Global diversity of enterococci and description of 18 novel species. bioRxiv.

Seemann, T. (2014). Prokka: rapid prokaryotic genome annotation. Bioinformatics, 30(14), 2068–2069.

The UniProt Consortium (2018). UniProt: a worldwide hub of protein knowledge. Nucleic Acids Research, 47(D1), D506–D515.

Törönen, P. et al. (2018). PANNZER2: a rapid functional annotation web server. Nucleic Acids Research, 46(W1), W84–W88.

van den Bent, I. et al. (2021). The power of universal contextualized protein embeddings in cross-species protein function prediction. Evolutionary Bioinformatics, 17.

van Kempen, M. et al. (2023). Foldseek: fast and accurate protein structure search. bioRxiv.

Xiong, X. et al. (2022). Emerging enterococcus pore-forming toxins with mhc/hla-i as receptors. Cell, 185(7), 1157–1171.e22.

Yao, S. et al. (2021). Netgo 2.0: improving large-scale protein function prediction with massive sequence, text, domain, family and network information. Nucleic acids research, 49(W1), W469–W475.

You, R. et al. (2018). GOLabeler: improving sequence-based large-scale protein function prediction by learning to rank. Bioinformatics, 34(14), 2465–2473.

Zhang, S. et al. (2018). Identification of a botulinum neurotoxin-like toxin in a commensal strain of enterococcus faecium. Cell Host & Microbe, 23(2), 169–176.e6.

Zhou, N. et al. (2019). The CAFA challenge reports improved protein function prediction and new functional annotations for hundreds of genes through experimental screens. Genome Biology, 20(1), 244.

