## Supplementary Material for "SAP: Synteny-aware gene function prediction for bacteria using protein embeddings"

### Table of Contents

**Table of Contents** 1
**Supplementary Text: Detailed examination of SAPdb** 2
Comparison of SAPdb operons to experimentally determined operons 2
Annotating SAPdb operons 5
Analysis of false positive enterococcal toxin predictions 7
**Supplementary Tables** 8
**Supplementary Figures** 19
**References** 24

### Supplementary Text: Detailed examination of SAPdb

#### Comparison of SAPdb operons to experimentally determined operons

The goal of constructing our own large-scale operon database (SAPdb) was to model the landscape of functions encapsulated within bacterial operons in sufficient detail to be able to leverage this resource to improve bacterial function prediction. In this section, we validate SAPdb as an adequate database of conserved operons by comparing it to a smaller operon database published previously titled Operon Database (ODB v4) (Okuda and Yoshizawa 2010).

In constructing SAPdb, we used information from ODB to refine thresholds such as operon length, number of genes in an operon, and intergenic distance between adjacent genes in an operon (Methods). ODB contains a curated list of experimentally determined operons obtained from the literature. The conserved operon database from ODB, referred to here as "ODB Conserved", is essentially an expansion on their known, experimentally determined operons, where the additional predicted operon instances were identified in additional genomes. Overall, our operon database was similar to the ODB Conserved database, in terms of operon length, number of genes in an operon and intergenic distance within operons (Fig S1-S3). Thus, with SAPdb we provide a database even more extensive than ODB Conserved, representing a more up-to-date and broader range of diversity within the bacterial kingdom.

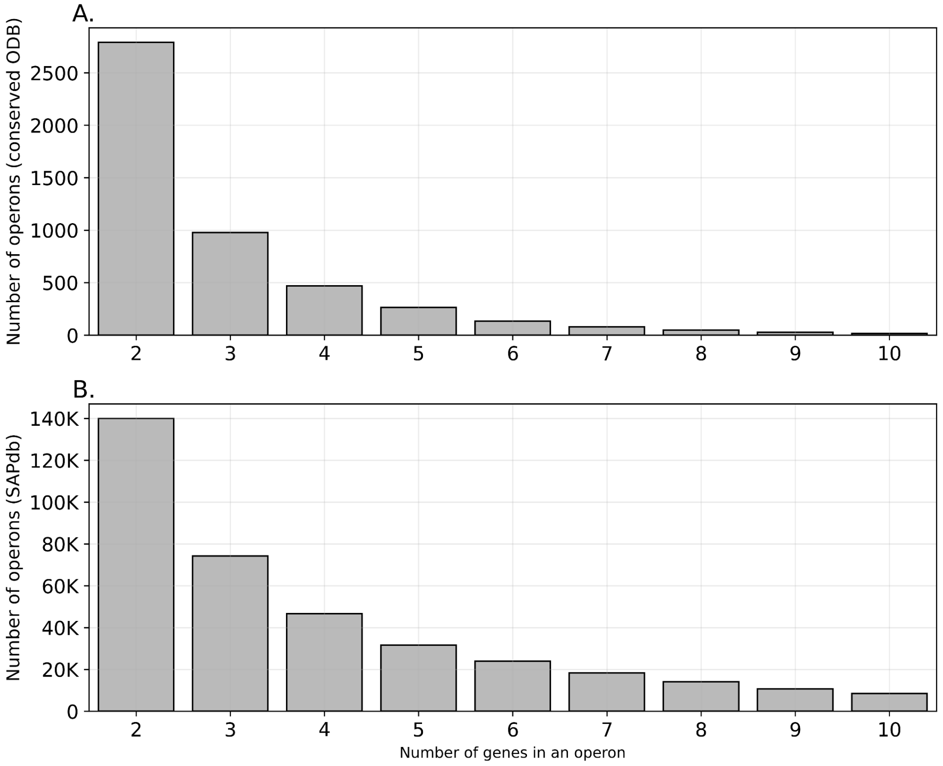

Fig S1. The operons in SAPdb have similar size distribution to those in ODB: A. operons in ODB. B. operons in SAPdb. Only non-singleton operons are shown.

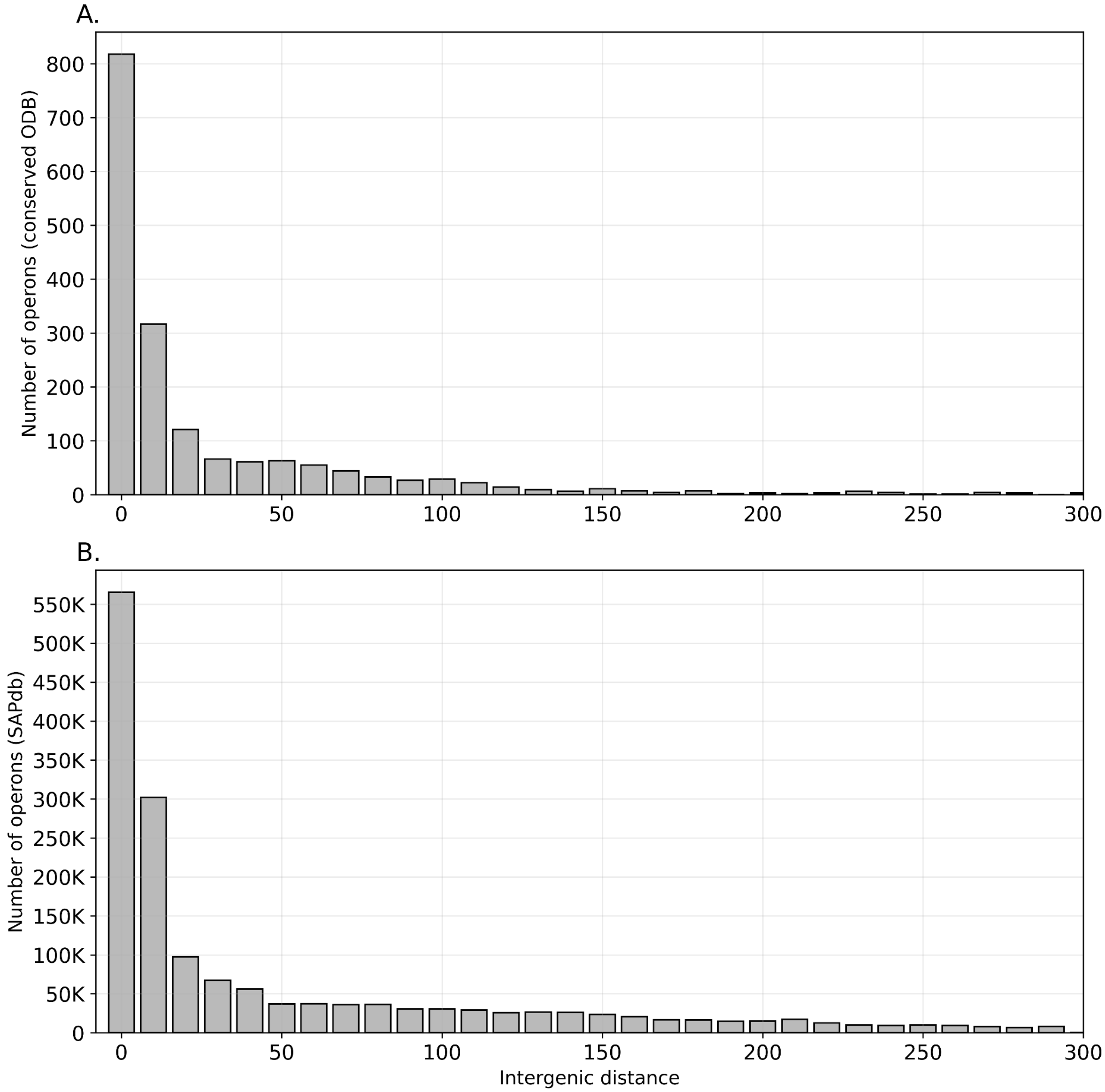


Fig S2. The operons in SAPdb have similar intergenic distance (bp) distribution to those in ODB: A. operons in ODB B. operons in SAPdb. Only non-singleton operons are shown.

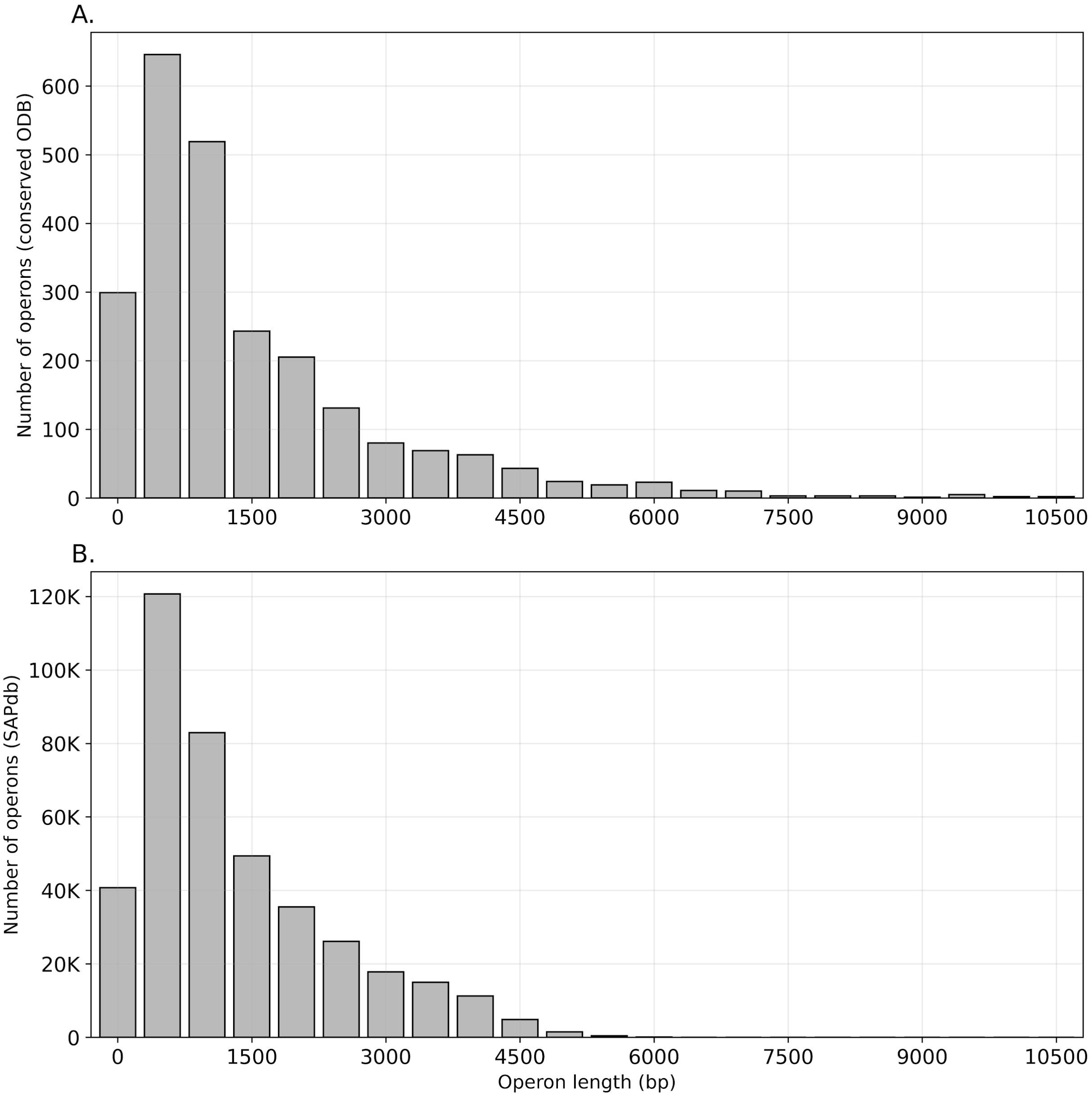


Fig S3. The operons in SAPdb have similar operon length distribution to those in ODB. A. operons in ODB. B. operons in SAPdb. Only non-singleton operons are shown.

With our approach, we also demonstrated that we could either partially or completely predict existing bacterial operons accurately. In order to assess whether we could reconstruct experimentally-determined operons in ODB for *E. coli*, based on our predicted operons in SAPdb, we extracted ODB operons containing at least one *E. coli* genes (2845 total operons, including 1071 non-singleton operons). We compared this to the predicted operons containing at least one *E. coli* genes  SAPdb, including 15,618 non-singleton operons, of which 87% (13,610 operons) partially or fully overlap with the known ODB operons. Full concordance is not expected, as our database is substantially larger and encompasses far more genes. In addition, our operon database is inherently redundant; a single experimentally known operon from ODB can be split into multiple predicted operons. Thus, it is highly likely that the remaining 2008 operons (23% of SAPdb) we predicted in addition to the ODB operons can be collapsed into a smaller number of actual operons.

We selected a few predictions showcasing the accuracy of SAPdb in reproducing experimentally known operons (Table S1). Although SAPdb often accurately reproduces known operons, it can sometimes capture additional genes in the flanking regions (i.e.IS3 family transposase genes flanking the relBEF toxin-antitoxin system in Table S1). Although this can give us additional information about conserved mechanisms related to the operon's mobilization, this can also lead to false positive annotation transfers using SAP.

Table S1. Our operon prediction algorithm can reproduce experimentally known operons in *E. coli*. Three example operons from SAPdb are shown, together with their corresponding operons in ODB, which were manually curated based on experimental studies.

| **Predicted operons in SAPdb** | | | | **Known operons in ODB** | | |
|---|---|---|---|---|---|---|
| **SAPdb Operon ID** | **Species** | ***E.coli gene*** | **Annotation** | **ODB Operon ID** | **Name** | **Definition** |
| 62671 | *Escherichia* , *Shigella*, *Citrobacter* | b4460 | L-arabinose transport system permease protein AraH | KO03244 | araFGH | High-affinity L-arabinose transport |
| | | b1900 | Arabinose import ATP-binding protein AraG (EC 7.5.2.12) | | | |
| | | b1901 | L-arabinose-binding periplasmic protein (ABP) | | | |
| 62674 | *Escherichia* , *Shigella*, *Bacillus* | b1879 | Flagellar biosynthetic protein FlhB | KO03228 | flhAB | Flagellar biosynthesis |
| | | b1880 | Flagellar biosynthesis protein FlhA | | | |
| 62755 | *Citrobacter*, *Enterobacter*, *Klebsiella*, *Leclercia*, *Escherichia*, | - | IS3 family transposase | - | - | |
| | | - | IS3 family transposase | - | - | |
| | | b1562 | Toxic protein HokD | KO03197 | relBEF | toxin-antitoxin system |
| | | b1563 | mRNA interferase toxin RelE (EC 3.1.-.-) (Endoribonuclease RelE) (Toxin RelE) | | | |
| | | b1564 | Antitoxin RelB | | | |

#### Annotating SAPdb operons

Although GTDB is the largest catalog of representative bacterial genomes, it does not catalog experimental annotations for the gene sequences. Thus, we assigned function to our predicted SAPdb operons by transferring GO terms from similar protein sequences in the SwissProt database, using thresholds on both the pairwise identity and significance (Methods). When transferring GO terms to SAPdb operons, we aimed to balance minimizing false positives while retaining as many annotations as possible, in order to avoid an operon set that was too sparsely annotated to be useful for predicting gene function. In our full SwissProt dataset experiments, 95.6% of the non-singleton operons (388,377 out of 406,293) were annotated with at least one

GO term; however, in remote homology experiments, SAPdb was a lot more sparse in annotations. Since we removed training proteins that had more than a certain predefined level of sequence homology to any of the test proteins in order to create the remote homology datasets, there were only rare proteins with no homologs left in the database. Given that such rare proteins are less likely to be studied, annotated or even carry out any function within the cell, this loss of functional information was expected. Since SAPdb is a major source of annotation that SAP relies on, SAP's prediction coverage also suffers from this loss of information, and we observed that the difference in coverage between SAP and DeepGOPlus widens as more annotations are lost (Table S13 and S14).

**Table S2.** GO term annotations in SAPdb were more sparse in the remote homology detection experiments. Percentage of entries in SAPdb that were annotated with at least one GO term for each GO term category and every organism in our SwissProt benchmarks.

| | **40** | **50** | **60** | **70** | **80** | **Full** |
|---|---|---|---|---|---|---|
| **Organism** | **Biological process** | | | | | |
| *E. coli* | 74.66% | 80.23% | 82.33% | 83.84% | 84.16% | 84.83% |
| *M. tuberculosis* | 80.32% | 83.55% | 84.65% | 85.07% | 85.21% | 85.18% |
| *B. subtilis* | 79.30% | 83.82% | 84.63% | 85.17% | 85.09% | 85.15% |
| *P. aeruginosa* | 79.21% | 82.10% | 82.79% | 83.89% | 84.34% | 85.16% |
| *S. typhimurium* | 83.56% | 84.44% | 84.80% | 84.96% | 85.03% | 85.16% |
| | **Molecular function** | | | | | |
| *E. coli* | 77.37% | 83.08% | 85.77% | 87.24% | 87.84% | 88.38% |
| *M. tuberculosis* | 83.08% | 86.40% | 87.57% | 88.13% | 88.38% | 88.59% |
| *B. subtilis* | 81.55% | 85.73% | 87.51% | 88.06% | 88.48% | 88.58% |
| *P. aeruginosa* | 82.03% | 84.94% | 86.35% | 87.47% | 87.97% | 88.59% |
| *S. typhimurium* | 87.27% | 87.90% | 88.24% | 88.45% | 88.49% | 88.59% |
| | **Cellular component** | | | | | |
| *E. coli* | 69.11% | 75.94% | 79.57% | 81.25% | 81.94% | 82.79% |
| *M. tuberculosis* | 76.02% | 80.00% | 81.83% | 82.49% | 82.82% | 83.06% |
| *B. subtilis* | 75.50% | 79.87% | 81.98% | 82.76% | 82.97% | 83.06% |
| *P. aeruginosa* | 75.71% | 78.92% | 80.86% | 81.92% | 82.44% | 83.05% |
| *S. typhimurium* | 81.37% | 82.26% | 82.61% | 82.84% | 82.91% | 83.06% |

Furthermore, we note that annotations were transferred at uneven rates for different categories of GO terms, consistent with previous findings in the literature. When sequence homology is used as the basis for transferring annotation, GO terms in the Molecular Function (MFO) category are more likely to be transferred than those for Biological Process (BPO) and Cellular Component (CCO), as MFO can more readily be modeled using the primary sequence, or features derived from it (Zhou et al. 2019). We also report that the number of operons annotated with at least one GO term is the largest for the MFO category, even though the total number of MFO terms available in the SwissProt dataset is significantly smaller than that of BPO (Table S3).

**Table S3.** Annotation statistics and information content (IC) of operons in SAPdb. GO terms in the Molecular Function (MFO) category are more likely to be transferred when using sequence homology for annotation transfer.

| Annotation statistic | BPO[a] | MFO[b] | CCO[c] |
|---|---|---|---|
| # of annotated operons (%) | 268,773 (70%) | 311,424 (80%) | 268,359 (70%) |
| Range of # of GO terms in an annotated operon | [1, 41] | [1, 17] | [1, 20] |
| Average # of GO terms per gene in an annotated operon | 0.556 | 0.543 | 0.456 |
| Average IC of GO terms in an annotated operon | 10.73 | 8.786 | 6.022 |
| Average IC of GO terms per gene in an annotated operon | 3.345 | 2.711 | 1.762 |
| Total # of GO terms in the SwissProt database | 16281 | 6308 | 2565 |

[a]Biological Process
[b]Molecular Function
[c]Cellular Component

#### Analysis of false positive enterococcal toxin predictions

**Table S4.** SAP associated 59 enterococcal proteins with the delta toxin operon from SAPdb (operon ID 395786), found in *Clostridium* and *Roseburia*. The contents of this operon are shown here.

| Operon | Cluster | Gene | Uniprot ID |
|---|---|---|---|
| 395786 | 75445122 | Heat-labile enterotoxin B chain | P01558 |
| | 84731674 | Uncharacterized lipoprotein YsaB | Q83J37 |

Forty-eight genes were associated with the SAPdb toxin operon, but did not have structural similarity to a known toxin protein fold.  The SAPdb toxin operon codes for two membrane-associated proteins:  1) the toxin gene itself, and 2) another membrane protein of unknown function. Based on FoldSeek search results, 33 of these 48 genes  appeared to perform other functions related to the cell membrane, involving signaling, secretion, and pore formation. To gain further insight into why these 48 additional genes without toxin structural folds were matched to the delta toxin operon, we compared the Euclidean similarity of embeddings vectors extracted from all 59 genes to the embedding vector of the known delta toxin operon (Table S4). While we did not find any significant differences in terms of the absolute similarity values between those 11 that had structural similarity to the delta toxin operon and the rest, we noted that the SAPdb toxin operon was consistently ranked in the top two among all SAPdb operons assigned to the 11 likely toxins, whereas for the remaining 48 genes this was not the case. This suggests that the ranking could be used in future versions of SAP as a proxy to infer confidence in operon assignments and to reduce false positives.

### Supplementary Tables

**Supplementary Table. S5.** Genome metadata and assembly statistics of the 19 *Enterococcus* genomes used in the *Enterococcus* diversity dataset

| Species | Clade[1] | NCBI Accession | Genome size (bp) | N50 (bp) | Number of components[2] | Completeness[3] | Contamination[4] | Number of CDS[5] |
|---|---|---|---|---|---|---|---|---|
| *E. caccae* | I | GCA_000407145.1 | 3551922 | 1036503 | 7 | 98.99 | 1.01 | 3244 |
| *E. haemoperoxidus* | I | GCA_000407165.1 | 3581919 | 2345301 | 2 | 100 | 1.52 | 3207 |
| *E. moraviensis* | I | GCA_000407445.1 | 3586110 | 878836 | 6 | 98.99 | 0 | 3337 |
| *E. durans* | II | GCA_000407265.1 | 3170574 | 972116 | 11 | 98.99 | 0 | 3031 |
| *E. hirae* | II | | 2882665 | 676112 | 6 | 98.99 | 0 | 2667 |
| *E. pernyi* | II | GCA_000407465.1 | 3078858 | 2163032 | 14 | 98.99 | 0 | 2891 |
| *E. phoeniculicola* | II | GCA_000407505.1 | 3910972 | 889011 | 11 | 100 | 0 | 3507 |
| *E. villorum* | II | GCA_000407205.1 | 3060724 | 1302785 | 4 | 98.99 | 0 | 2821 |
| *E. avium* | III | GCA_000407245.1 | 4614282 | 652010 | 13 | 98.99 | 0 | 4504 |
| *E. gilvus* | III | GCA_000407545.1 | 4179913 | 3038269 | 5 | 98.99 | 0 | 4111 |
| *E. malodoratus* | III | GCA_000407185.1 | 4627134 | 1188036 | 7 | 98.99 | 0 | 4500 |
| *E. pallens* | III | GCA_000407485.1 | 5437454 | 1015307 | 13 | 100 | 0 | 5196 |
| *E. raffinosus* | III | GCA_000407525.1 | 4310366 | 1007288 | 9 | 98.99 | 0 | 4202 |
| *E. asini* | IV | GCA_000407365.1 | 2572706 | 2038454 | 2 | 98.99 | 0 | 2429 |
| *E. cecorum* | IV | GCA_000492155.1 | 2408477 | 1060610 | 6 | 98.99 | 0 | 2375 |
| *E. columbae* | IV | GCA_000407225.1 | 2545099 | 421089 | 9 | 98.99 | 0 | 2327 |
| *E. dispar* | IV | GCA_000407585.1 | 2812918 | 2801753 | 4 | 98.99 | 0 | 2635 |
| *E. saccharolyticus* | IV | GCA_000407285.1 | 2604038 | 2432614 | 2 | 98.99 | 0 | 2586 |
| *E. sulfureus* | IV | GCA_000407605.1 | 2301651 | 733717 | 8 | 98.99 | 0 | 2176 |

[1]*Enterococcus* clades, defined by (Lebreton et al., 2017).

[2]Number of contigs/scaffolds or chromosomes.

[3,4]Calculated using CheckM (Parks et al., 2015).

[5]Total CDS predicted by Prokka (Seemann, 2014).

**Supplementary Table. S6.** $S_{min}$ values for the full SwissProt dataset.

| $S_{min}$ values for each of five species[1,2] | | | | | |
|---|---|---|---|---|---|
| **Method** | ***EC*** | ***MT*** | ***BS*** | ***PA*** | ***ST*** |
| **Biological process** | | | | | |
| BLAST | 20.97 | 22.93 | 15.99 | 19.02 | 8.53 |
| Pfam | 108.50 | 161.14 | 126.22 | 122.86 | 129.13 |
| knn (T5) | 18.61 | 18.44 | 9.84 | 12.76 | 8.83 |
| knn | 17.44 | 18.24 | 9.02 | 12.18 | 8.37 |
| SAP-operon | **7.10** | **7.60** | **3.27** | **4.41** | 4.90 |
| SAP | **7.10** | **7.60** | **3.27** | **4.41** | 4.90 |
| DeepGOPlus | 16.45 | 14.94 | 5.68 | 10.39 | **2.85** |
| **Molecular function** | | | | | |
| BLAST | 10.97 | 11.07 | 7.99 | 8.25 | 5.51 |
| Pfam | 30.87 | 52.74 | 35.04 | 35.48 | 37.17 |
| knn (T5) | 10.01 | 8.99 | 4.74 | 4.52 | 5.72 |
| knn | 9.12 | 8.04 | 4.77 | 4.49 | 5.67 |
| SAP-operon | **3.63** | **3.51** | **1.88** | **1.66** | 2.72 |
| SAP | **3.63** | **3.51** | **1.88** | **1.66** | 2.72 |
| DeepGOPlus | 8.68 | 6.36 | 2.57 | 2.67 | **1.68** |
| **Cellular component** | | | | | |
| BLAST | 6.46 | 5.74 | 3.87 | 4.17 | 2.53 |
| Pfam | 30.29 | 35.91 | 34.37 | 31.40 | 32.15 |
| knn (T5) | 4.87 | 4.99 | 1.76 | 1.81 | 2.37 |
| knn | 5.02 | 5.00 | 1.84 | 2.05 | 2.29 |
| SAP-operon | **1.64** | **1.85** | **0.49** | **0.82** | 1.35 |
| SAP | **1.64** | **1.85** | **0.49** | **0.82** | 1.35 |
| DeepGOPlus | 4.92 | 4.63 | 1.55 | 1.77 | **1.18** |

[1] EC: *Escherichia coli*, MT: *Mycobacterium tuberculosis*, BS: *Bacillus subtilis*, PA: *Pseudomonas aeruginosa* and ST: *Salmonella typhimurium*.

[2] Lowest value for each species and for each GO category are shown in bold.

**Supplementary Table. S7.** Area under the precision/recall curve for the full SwissProt dataset, for all 5 bacterial organisms and three GO categories.

| Area under precision/recall curve for each of five species[1,2] | | | | | |
|---|---|---|---|---|---|
| **Method** | ***EC*** | ***MT*** | ***BS*** | ***PA*** | ***ST*** |
| **Biological process** | | | | | |
| BLAST | 0.519 | 0.497 | 0.686 | 0.642 | 0.642 |
| Pfam | 0.648 | 0.556 | 0.582 | 0.623 | 0.623 |
| knn (T5) | 0.529 | 0.514 | 0.76 | 0.714 | 0.714 |
| knn | 0.544 | 0.533 | 0.80 | 0.714 | 0.714 |
| DeepGOPlus | 0.564 | 0.600 | 0.862 | 0.797 | 0.797 |
| SAP-operon | **0.855** | **0.805** | **0.907** | **0.908** | **0.908** |
| SAP | **0.855** | **0.805** | **0.907** | **0.908** | **0.908** |
| **Molecular function** | | | | | |
| BLAST | 0.569 | 0.573 | 0.658 | 0.644 | 0.644 |
| Pfam | 0.686 | 0.599 | 0.571 | 0.595 | 0.595 |
| knn (T5) | 0.523 | 0.602 | 0.764 | 0.774 | 0.774 |
| knn | 0.560 | 0.640 | 0.790 | 0.780 | 0.780 |
| DeepGOPlus | 0.613 | 0.719 | 0.857 | 0.899 | 0.899 |
| SAP-operon | **0.872** | **0.849** | **0.887** | **0.921** | **0.921** |
| SAP | **0.872** | **0.849** | **0.887** | **0.921** | **0.921** |
| **Cellular component** | | | | | |
| BLAST | 0.541 | 0.324 | 0.701 | 0.724 | 0.724 |
| Pfam | 0.675 | 0.598 | 0.608 | 0.634 | 0.634 |
| knn (T5) | 0.641 | 0.344 | 0.858 | 0.849 | 0.849 |
| knn | 0.633 | 0.335 | 0.879 | 0.853 | 0.853 |
| DeepGOPlus | 0.747 | 0.512 | 0.931 | 0.917 | 0.917 |
| SAP-operon | **0.919** | **0.831** | **0.954** | **0.945** | **0.945** |
| SAP | **0.919** | **0.831** | **0.954** | **0.945** | **0.945** |

[1] EC: *Escherichia coli*, MT: *Mycobacterium tuberculosis*, BS: *Bacillus subtilis*, PA: *Pseudomonas aeruginosa* and ST: *Salmonella typhimurium*

[2] The highest value for each species and for each GO category are shown in bold

**Supplementary Table. S8.** $F_{max}$ values for the entire *E. coli* SwissProt benchmark sets.

| Method | 40 | 50 | 60 | 70 | 80 | Full |
|---|---|---|---|---|---|---|
| **Biological process** | | | | | | |
| BLAST | 0.505 | 0.526 | 0.540 | 0.556 | 0.560 | 0.570 |
| Pfam | 0.610 | 0.610 | 0.610 | 0.610 | 0.610 | 0.610 |
| knn (T5) | 0.464 | 0.510 | 0.574 | 0.596 | 0.607 | 0.625 |
| knn | 0.536 | 0.584 | 0.623 | 0.635 | 0.639 | 0.646 |
| DeepGOPlus | 0.568 | 0.601 | 0.624 | 0.636 | 0.644 | 0.648 |
| SAP-operon | 0.720 | 0.792 | 0.824 | 0.840 | 0.858 | 0.877 |
| SAP | 0.724 | 0.800 | 0.830 | 0.846 | 0.865 | 0.877 |
| **Molecular function** | | | | | | |
| BLAST | 0.554 | 0.573 | 0.585 | 0.599 | 0.603 | 0.613 |
| Pfam | 0.650 | 0.650 | 0.650 | 0.650 | 0.650 | 0.650 |
| knn (T5) | 0.475 | 0.528 | 0.595 | 0.611 | 0.618 | 0.632 |
| knn | 0.558 | 0.598 | 0.656 | 0.666 | 0.669 | 0.675 |
| DeepGOPlus | 0.605 | 0.641 | 0.649 | 0.674 | 0.680 | 0.686 |
| SAP-operon | 0.725 | 0.803 | 0.833 | 0.850 | 0.865 | 0.886 |
| SAP | 0.739 | 0.810 | 0.840 | 0.855 | 0.872 | 0.886 |
| **Cellular component** | | | | | | |
| BLAST | 0.502 | 0.519 | 0.532 | 0.546 | 0.554 | 0.569 |
| Pfam | 0.625 | 0.625 | 0.625 | 0.625 | 0.625 | 0.625 |
| knn (T5) | 0.560 | 0.610 | 0.691 | 0.712 | 0.732 | 0.738 |
| knn | 0.610 | 0.650 | 0.707 | 0.715 | 0.723 | 0.731 |
| DeepGOPlus | 0.645 | 0.705 | 0.721 | 0.706 | 0.726 | 0.745 |
| SAP-operon | 0.769 | 0.835 | 0.866 | 0.887 | 0.905 | 0.925 |
| SAP | 0.774 | 0.842 | 0.873 | 0.893 | 0.911 | 0.925 |

**Supplementary Table. S9.** $F_{max}$ values for the entire *M. tuberculosis* SwissProt benchmark sets.

| Method | 40 | 50 | 60 | 70 | 80 | Full |
|---|---|---|---|---|---|---|
| **Biological process** | | | | | | |
| BLAST | 0.525 | 0.521 | 0.531 | 0.532 | 0.539 | 0.543 |
| Pfam | 0.513 | 0.513 | 0.513 | 0.513 | 0.513 | 0.513 |
| knn (T5) | 0.520 | 0.597 | 0.613 | 0.615 | 0.618 | 0.618 |
| knn | 0.575 | 0.617 | 0.629 | 0.630 | 0.633 | 0.636 |
| DeepGOPlus | 0.627 | 0.645 | 0.670 | 0.667 | 0.667 | 0.669 |
| SAP-operon | 0.698 | 0.750 | 0.786 | 0.812 | 0.825 | 0.838 |
| SAP | 0.703 | 0.750 | 0.787 | 0.813 | 0.828 | 0.838 |
| **Molecular function** | | | | | | |
| BLAST | 0.589 | 0.582 | 0.580 | 0.584 | 0.591 | 0.593 |
| Pfam | 0.549 | 0.549 | 0.549 | 0.549 | 0.549 | 0.549 |
| knn (T5) | 0.586 | 0.654 | 0.672 | 0.677 | 0.683 | 0.681 |
| knn | 0.654 | 0.706 | 0.714 | 0.718 | 0.722 | 0.723 |
| DeepGOPlus | 0.709 | 0.736 | 0.745 | 0.752 | 0.750 | 0.755 |
| SAP-operon | 0.748 | 0.800 | 0.830 | 0.851 | 0.864 | 0.869 |
| SAP | 0.748 | 0.801 | 0.832 | 0.856 | 0.866 | 0.869 |
| **Cellular component** | | | | | | |
| BLAST | 0.401 | 0.397 | 0.394 | 0.396 | 0.396 | 0.397 |
| Pfam | 0.541 | 0.541 | 0.541 | 0.541 | 0.541 | 0.541 |
| knn (T5) | 0.434 | 0.513 | 0.520 | 0.517 | 0.510 | 0.507 |
| knn | 0.431 | 0.504 | 0.505 | 0.505 | 0.502 | 0.500 |
| DeepGOPlus | 0.570 | 0.577 | 0.575 | 0.573 | 0.572 | 0.567 |
| SAP-operon | 0.634 | 0.700 | 0.753 | 0.804 | 0.835 | 0.846 |
| SAP | 0.638 | 0.704 | 0.756 | 0.807 | 0.835 | 0.846 |

**Supplementary Table. S10.** $F_{max}$ values for the entire *B. subtilis* SwissProt benchmark sets.

| Method | 40 | 50 | 60 | 70 | 80 | Full |
|---|---|---|---|---|---|---|
| **Biological process** | | | | | | |
| BLAST | 0.628 | 0.636 | 0.634 | 0.638 | 0.638 | 0.639 |
| Pfam | 0.582 | 0.582 | 0.582 | 0.582 | 0.582 | 0.582 |
| knn (T5) | 0.700 | 0.768 | 0.783 | 0.788 | 0.790 | 0.791 |
| knn | 0.752 | 0.809 | 0.821 | 0.826 | 0.827 | 0.828 |
| DeepGOPlus | 0.791 | 0.835 | 0.841 | 0.852 | 0.857 | 0.857 |
| SAP-operon | 0.822 | 0.881 | 0.897 | 0.907 | 0.913 | 0.921 |
| SAP | 0.828 | 0.885 | 0.900 | 0.909 | 0.914 | 0.921 |
| **Molecular function** | | | | | | |
| BLAST | 0.622 | 0.630 | 0.623 | 0.624 | 0.625 | 0.625 |
| Pfam | 0.571 | 0.571 | 0.571 | 0.571 | 0.571 | 0.571 |
| knn (T5) | 0.672 | 0.758 | 0.782 | 0.789 | 0.791 | 0.791 |
| knn | 0.735 | 0.789 | 0.805 | 0.809 | 0.813 | 0.814 |
| DeepGOPlus | 0.775 | 0.830 | 0.836 | 0.842 | 0.850 | 0.841 |
| SAP-operon | 0.798 | 0.860 | 0.878 | 0.888 | 0.891 | 0.898 |
| SAP | 0.803 | 0.864 | 0.884 | 0.892 | 0.893 | 0.898 |
| **Cellular component** | | | | | | |
| BLAST | 0.622 | 0.623 | 0.631 | 0.637 | 0.638 | 0.638 |
| Pfam | 0.608 | 0.608 | 0.608 | 0.608 | 0.608 | 0.608 |
| knn (T5) | 0.766 | 0.859 | 0.874 | 0.883 | 0.883 | 0.884 |
| knn | 0.811 | 0.870 | 0.888 | 0.896 | 0.897 | 0.898 |
| DeepGOPlus | 0.820 | 0.848 | 0.863 | 0.888 | 0.864 | 0.885 |
| SAP-operon | 0.878 | 0.920 | 0.926 | 0.935 | 0.941 | 0.944 |
| SAP | 0.874 | 0.925 | 0.929 | 0.939 | 0.941 | 0.944 |

**Supplementary Table. S11.** $F_{max}$ values for the entire *P. aeruginosa* SwissProt benchmark sets.

| **Method** | **40** | **50** | **60** | **70** | **80** | **Full** |
|---|---|---|---|---|---|---|
| **Biological process** | | | | | | |
| BLAST | 0.650 | 0.666 | 0.680 | 0.686 | 0.684 | 0.683 |
| Pfam | 0.579 | 0.579 | 0.579 | 0.579 | 0.579 | 0.579 |
| knn (T5) | 0.629 | 0.731 | 0.786 | 0.798 | 0.797 | 0.796 |
| knn | 0.681 | 0.769 | 0.794 | 0.799 | 0.797 | 0.797 |
| DeepGOPlus | 0.749 | 0.785 | 0.812 | 0.820 | 0.816 | 0.824 |
| SAP-operon | 0.754 | 0.835 | 0.879 | 0.892 | 0.916 | 0.927 |
| SAP | 0.755 | 0.837 | 0.878 | 0.896 | 0.922 | 0.927 |
| **Molecular function** | | | | | | |
| BLAST | 0.712 | 0.721 | 0.716 | 0.708 | 0.702 | 0.699 |
| Pfam | 0.534 | 0.534 | 0.534 | 0.534 | 0.534 | 0.534 |
| knn (T5) | 0.679 | 0.801 | 0.848 | 0.858 | 0.856 | 0.853 |
| knn | 0.765 | 0.849 | 0.863 | 0.863 | 0.857 | 0.854 |
| DeepGOPlus | 0.810 | 0.856 | 0.883 | 0.894 | 0.882 | 0.883 |
| SAP-operon | 0.789 | 0.879 | 0.913 | 0.921 | 0.932 | 0.938 |
| SAP | 0.784 | 0.882 | 0.915 | 0.925 | 0.939 | 0.938 |
| **Cellular component** | | | | | | |
| BLAST | 0.658 | 0.687 | 0.692 | 0.701 | 0.701 | 0.700 |
| Pfam | 0.560 | 0.560 | 0.560 | 0.560 | 0.560 | 0.560 |
| knn (T5) | 0.743 | 0.838 | 0.876 | 0.894 | 0.900 | 0.898 |
| knn | 0.780 | 0.868 | 0.894 | 0.900 | 0.900 | 0.900 |
| DeepGOPlus | 0.819 | 0.847 | 0.874 | 0.877 | 0.900 | 0.887 |
| SAP-operon | 0.823 | 0.890 | 0.914 | 0.927 | 0.934 | 0.945 |
| SAP | 0.830 | 0.893 | 0.914 | 0.931 | 0.943 | 0.945 |

**Supplementary Table. S12.** $F_{max}$ values for the entire *S. typhimurium* SwissProt benchmark sets.

| Method | 40 | 50 | 60 | 70 | 80 | Full |
|---|---|---|---|---|---|---|
| **Biological process** | | | | | | |
| BLAST | 0.675 | 0.728 | 0.775 | 0.800 | 0.822 | 0.852 |
| Pfam | 0.579 | 0.579 | 0.579 | 0.579 | 0.579 | 0.579 |
| knn (T5) | 0.634 | 0.739 | 0.795 | 0.848 | 0.875 | 0.869 |
| knn | 0.774 | 0.855 | 0.889 | 0.910 | 0.919 | 0.880 |
| DeepGOPlus | 0.811 | 0.860 | 0.896 | 0.911 | 0.922 | 0.928 |
| SAP-operon | 0.811 | 0.871 | 0.894 | 0.906 | 0.908 | 0.905 |
| SAP | 0.811 | 0.871 | 0.895 | 0.907 | 0.910 | 0.905 |
| **Molecular function** | | | | | | |
| BLAST | 0.674 | 0.723 | 0.751 | 0.782 | 0.811 | 0.814 |
| Pfam | 0.559 | 0.559 | 0.559 | 0.559 | 0.559 | 0.559 |
| knn (T5) | 0.589 | 0.694 | 0.753 | 0.818 | 0.856 | 0.829 |
| knn | 0.761 | 0.844 | 0.868 | 0.894 | 0.904 | 0.837 |
| DeepGOPlus | 0.800 | 0.845 | 0.875 | 0.892 | 0.904 | 0.911 |
| SAP-operon | 0.796 | 0.863 | 0.884 | 0.893 | 0.889 | 0.883 |
| SAP | 0.798 | 0.862 | 0.885 | 0.895 | 0.890 | 0.883 |
| **Cellular component** | | | | | | |
| BLAST | 0.644 | 0.705 | 0.737 | 0.774 | 0.815 | 0.871 |
| Pfam | 0.616 | 0.616 | 0.616 | 0.616 | 0.616 | 0.616 |
| knn (T5) | 0.794 | 0.848 | 0.871 | 0.898 | 0.918 | 0.916 |
| knn | 0.824 | 0.881 | 0.909 | 0.923 | 0.941 | 0.917 |
| DeepGOPlus | 0.819 | 0.868 | 0.898 | 0.901 | 0.917 | 0.936 |
| SAP-operon | 0.856 | 0.896 | 0.912 | 0.920 | 0.922 | 0.918 |
| SAP | 0.858 | 0.899 | 0.916 | 0.922 | 0.923 | 0.918 |

**Supplementary Table. S13.** Prediction coverage[1] (in %) for the full SwissProt dataset.

| Prediction coverage for each of five species (%)[2] | | | | | |
|---|---|---|---|---|---|
| **Method** | ***EC*** | ***MT*** | ***BS*** | ***PA*** | ***ST*** |
| **Biological process** | | | | | |
| BLAST | 65.66 | 66.14 | 75.58 | 73.11 | 87.61 |
| Pfam | 81.41 | 71.19 | 80.69 | 75.03 | 84.42 |
| knn (T5) | 84.78 | 91.40 | 91.58 | 91.00 | 97.15 |
| knn | 88.29 | 90.16 | 94.88 | 94.12 | 97.82 |
| DeepGOPlus | 88.79 | 81.92 | 94.80 | 95.80 | 97.49 |
| SAP-operon | 92.48 | 82.36 | 90.68 | 92.32 | 90.45 |
| SAP | 92.66 | 82.36 | 90.68 | 92.32 | 90.45 |
| **Molecular function** | | | | | |
| BLAST | 74.04 | 69.51 | 75.25 | 81.36 | 89.23 |
| Pfam | 88.46 | 88.80 | 87.28 | 88.82 | 87.44 |
| knn (T5) | 89.64 | 96.53 | 91.47 | 98.59 | 97.06 |
| knn | 88.35 | 96.00 | 93.56 | 98.33 | 97.23 |
| DeepGOPlus | 98.05 | 92.80 | 89.69 | 94.86 | 96.25 |
| SAP-operon | 92.85 | 92.98 | 88.29 | 93.06 | 88.42 |
| SAP | 93.33 | 92.98 | 88.29 | 93.06 | 88.42 |
| **Cellular component** | | | | | |
| BLAST | 60.29 | 58.41 | 68.377 | 77.51 | 87.05 |
| Pfam | 81.65 | 76.35 | 81.981 | 84.43 | 89.68 |
| knn (T5) | 82.39 | 67.22 | 93.437 | 93.60 | 96.15 |
| knn | 81.36 | 64.92 | 94.153 | 93.43 | 96.76 |
| DeepGOPlus | 89.86 | 95.08 | 97.017 | 94.29 | 97.98 |
| SAP-operon | 93.32 | 92.86 | 95.943 | 91.35 | 96.96 |
| SAP | 93.53 | 92.86 | 95.943 | 91.35 | 96.96 |

[1] The number of test proteins annotated with at least one GO term at the threshold which maximizes the F1-score.

[2] EC: *Escherichia coli*, MT: *Mycobacterium tuberculosis*, BS: *Bacillus subtilis*, PA: *Pseudomonas aeruginosa* and ST: *Salmonella typhimurium*

**Supplementary Table. S14.** Prediction coverage[1] (in %) for the entire SwissProt benchmark evaluation, averaged over 5 bacterial organisms for three GO categories. The standard deviation across the 5 bacterial organisms is indicated in parentheses.

| Method | Maximum sequence % ID between the training and the test set | | | | | |
|---|---|---|---|---|---|---|
| | **40%** | **50%** | **60%** | **70%** | **80%** | **Full** |
| **Biological process** | | | | | | |
| BLAST | 71.16% (3.30%) | 70.20% (5.82%) | 70.21% (5.00%) | 71.76% (5.62%) | 72.02% (4.49%) | 73.62% (7.99%) |
| Pfam[2] | 78.55% (4.77%) | 78.55% (4.77%) | 78.55% (4.77%) | 78.55% (4.77%) | 78.55% (4.77%) | 78.55% (4.77%) |
| knn (T5) | 85.71% (4.99%) | 88.23% (6.43%) | 90.09% (3.74%) | 89.03% (4.80%) | 89.93% (4.49%) | 91.18% (3.92%) |
| knn | 85.33% (4.07%) | 91.58% (6.72%) | 93.23% (3.52%) | 92.40% (4.24%) | 92.76% (3.78%) | 93.06% (3.41%) |
| DeepGOPlus | 95.62% (1.56%) | 94.48% (3.45%) | 93.84% (4.99%) | 94.44% (3.16%) | 90.25% (6.16%) | 91.76% (5.73%) |
| SAP-operon | 85.48% (3.18%) | 87.36% (1.05%) | 87.93% (3.90%) | 89.95% (4.43%) | 91.42% (0.83%) | 89.66% (3.74%) |
| SAP | 84.04% (1.82%) | 87.93% (0.92%) | 88.11% (3.90%) | 89.17% (3.60%) | 91.73% (0.91%) | 89.69% (3.77%) |
| **Molecular function** | | | | | | |
| BLAST | 75.42% (3.39%) | 75.52% (2.93%) | 74.33% (3.73%) | 74.51% (4.34%) | 75.17% (4.20%) | 77.88% (6.82%) |
| Pfam[2] | 88.16% (0.67%) | 88.16% (0.67%) | 88.16% (0.67%) | 88.16% (0.67%) | 88.16% (0.67%) | 88.16% (0.67%) |
| knn (T5) | 89.55% (4.20%) | 92.16% (6.19%) | 94.44% (3.49%) | 94.46% (3.41%) | 94.62% (3.10%) | 94.66% (3.47%) |
| knn | 89.67% (5.32%) | 92.59% (6.82%) | 94.83% (3.31%) | 94.68% (3.32%) | 94.69% (3.49%) | 94.69% (3.55%) |
| DeepGOPlus | 93.79% (2.32%) | 93.98% (3.29%) | 94.46% (2.71%) | 93.98% (3.00%) | 95.25% (2.48%) | 94.33% (2.89%) |
| SAP-operon | 87.07% (3.14%) | 88.57% (1.60%) | 90.65% (1.65%) | 91.35% (1.97%) | 91.36% (2.33%) | 91.12% (2.26%) |
| SAP | 85.37% (2.80%) | 89.20% (1.75%) | 90.96% (1.49%) | 91.69% (2.02%) | 91.79% (2.50%) | 91.21% (2.34%) |
| **Cellular component** | | | | | | |
| BLAST | 71.81% (8.69%) | 67.46% (5.39%) | 68.98% (7.05%) | 70.76% (7.77%) | 67.96% (7.00%) | 70.33% (10.75%) |
| Pfam[2] | 82.82% (4.33%) | 82.82% (4.33%) | 82.82% (4.33%) | 82.82% (4.33%) | 82.82% (4.33%) | 82.82% (4.33%) |
| knn (T5) | 71.44% (12.05%) | 82.58% (11.93%) | 86.49% (8.77%) | 86.49% (9.561%) | 86.47% (9.99%) | 86.56% (10.77%) |
| knn | 74.37% (12.89%) | 83.43% (12.24%) | 86.28% (11.03%) | 86.31% (11.50%) | 86.56% (11.99%) | 86.12% (11.86%) |
| DeepGOPlus | 94.66% (1.63%) | 95.86% (1.52%) | 93.86% (1.86%) | 95.23% (1.89%) | 94.88% (2.90%) | 94.84% (2.82%) |
| SAP-operon | 85.93% (4.14%) | 91.17% (3.14%) | 90.54% (5.40%) | 91.42% (3.94%) | 93.50% (2.48%) | 94.09% (2.07%) |
| SAP | 85.87% (5.42%) | 91.01% (3.82%) | 92.08% (3.44%) | 92.91% (4.08%) | 93.89% (2.17%) | 94.13% (2.05%) |

[1] The number of test proteins annotated with at least one GO term at the threshold which maximizes the F1-score.

[2] Because the Pfam method uses a different training set for predictions, it is not possible to compare the change in the coverage across different train/test pairs.

**Supplementary Table S15.** Maximum % sequence identity of *Enterococcus* toxin genes to the 11 putative novel toxins predicted by SAP.

| epx gene type | epx gene locus tag | predicted gene locus tag | maximum identity (%) |
|---|---|---|---|
| epx1 | GCA_015300525.1_02780 | Ente_haem_BAA-382_V2_01251 | 38.89 |
| epx1 | GCA_002945555.1_00127 | Ente_haem_BAA-382_V2_01826 | 25.76 |
| epx1 | GCA_003319525.1_02660 | Ente_haem_BAA-382_V2_01826 | 25.76 |
| epx1 | GCA_004120285.1_01193 | Ente_haem_BAA-382_V2_01826 | 25.76 |
| epx1 | GCA_004125665.1_01702 | Ente_haem_BAA-382_V2_01826 | 25.76 |
| epx1 | GCA_004125915.1_01060 | Ente_haem_BAA-382_V2_01826 | 25.76 |
| epx1 | GCA_004125935.1_01411 | Ente_haem_BAA-382_V2_01826 | 25.76 |
| epx1 | GCA_004125945.1_01477 | Ente_haem_BAA-382_V2_01826 | 25.76 |
| epx1 | GCA_004125955.1_00929 | Ente_haem_BAA-382_V2_01826 | 25.76 |
| epx1 | GCA_004126025.1_01270 | Ente_haem_BAA-382_V2_01826 | 25.76 |
| epx1 | GCA_005236595.1_02731 | Ente_haem_BAA-382_V2_01826 | 25.76 |
| epx1 | GCA_005237765.1_02590 | Ente_haem_BAA-382_V2_01826 | 25.76 |
| epx1 | GCA_005238535.1_01041 | Ente_haem_BAA-382_V2_01826 | 25.76 |
| epx1 | GCA_015302065.1_02752 | Ente_haem_BAA-382_V2_01826 | 25.76 |
| epx1 | GCA_015333445.1_02601 | Ente_haem_BAA-382_V2_01826 | 25.76 |
| epx1 | GCA_015335045.1_02694 | Ente_haem_BAA-382_V2_01826 | 25.76 |
| epx1 | GCA_015336125.1_02821 | Ente_haem_BAA-382_V2_01826 | 25.76 |
| epx1 | GCA_015338255.1_02588 | Ente_haem_BAA-382_V2_01826 | 25.76 |
| epx4 | GCA_017356565.1_01452 | Ente_mund_ATCC882_V5_00469 | 60.00 |
| epx4 | GCA_015506475.1_02489 | Ente_haem_BAA-382_V2_01251 | 28.85 |
| epx4 | GCA_015507715.1_02409 | Ente_haem_BAA-382_V2_01251 | 28.85 |
| epx4 | GCA_018089265.1_02226 | Ente_haem_BAA-382_V2_01251 | 28.85 |

### Supplementary Figures

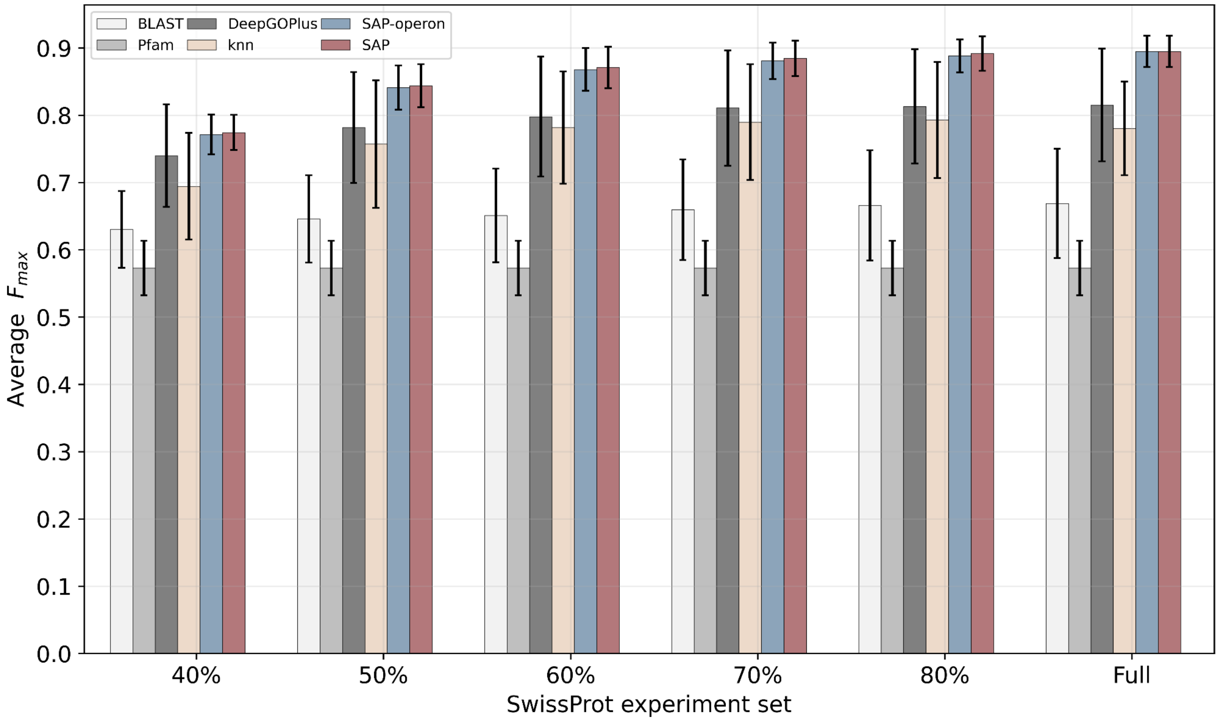


**Supplementary Fig. S4.** average $F_{max}$ values in Molecular Function Ontology (MFO), across benchmarking datasets for five bacteria in our experimental setup, and the error bars show the corresponding standard deviation of each method, across five species. Note that bar plots for the Pfam baseline are identical for all 6 experiment sets because Pfam uses a different training set, independent of our experimental design.

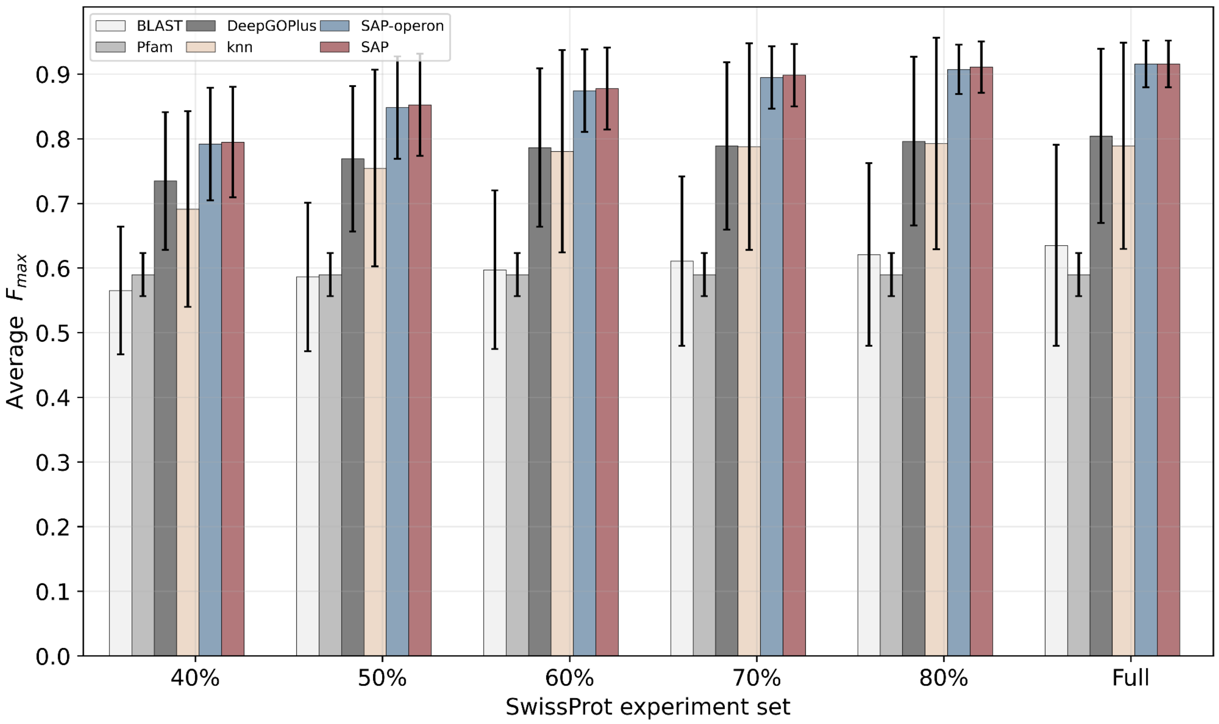


**Supplementary Fig. S5.** average $F_{max}$ values in Cellular Component Ontology (CCO), across benchmarking datasets for five bacteria in our experimental setup, and the error bars show the corresponding standard deviation of each method, across five species. Note that bar plots for the Pfam baseline are identical for all 6 experiment sets because Pfam uses a different training set, independent of our experimental design.

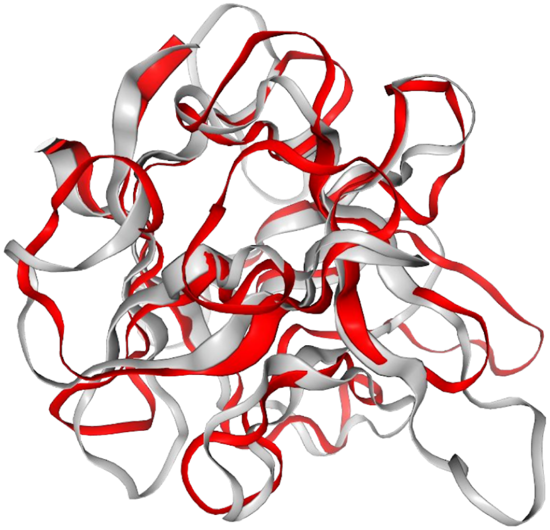

Sequence ID: 0.058
TM-Score: 0.76

Query
*E. haemoperoxidus* BAA-382
0174

```
38 NRNYVIKANPEGFKAKYWKTRYVD----VGSDWVNLTTDKDN-GSSYQLVTNGIVQENVAYKIYDSGSG-KYLNYQAGNT
        +  +++                    +     V+    VN  +    +     + L            +     +  + L  +A
```

Target
*epx1*

```
1 MAHVTLQSLS--NNDLCLDVYGENGDKTVAGGSVNGWSCHGSWNQVWGLDK--------EERYRSRVASDRCLTVNAD--
```

**Supplementary Fig. S6.** SAP predicts a possible new variant of pore-forming toxin gene in *E. haemoperoxidus*, distantly related to the known toxin Epx1, as evidenced by the highly significant structural alignment to the known toxin Epx1. Gene 0174 (red ribbon, query) had a TM-alignment score of 0.76 when we aligned its structure to Epx1 with Foldseek (gray ribbon, target), despite low sequence identity (5.8%). The protein's sequence alignment is shown below the 3D structural alignment.

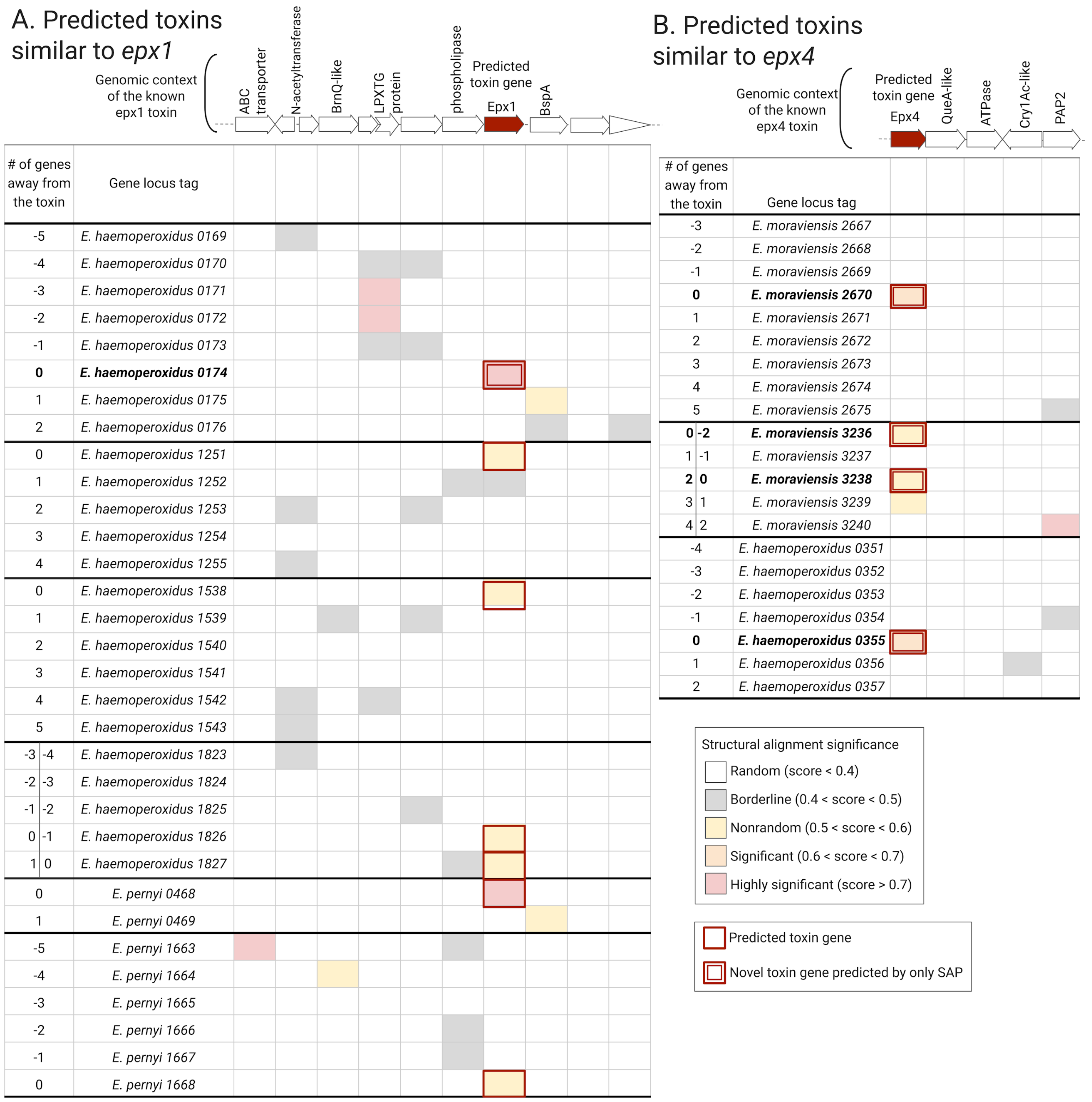
A. Predicted toxins similar to epx1
Genomic context of the known epx1 toxin
ABC transporter
N-acetyltransferase
BrnQ-like
LPXTG protein
phospholipase
Predicted toxin gene
Epx1
BspA
### of genes away from the toxin
Gene locus tag
-5 E. haemoperoxidus 0169
-4 E. haemoperoxidus 0170
-3 E. haemoperoxidus 0171
-2 E. haemoperoxidus 0172
-1 E. haemoperoxidus 0173
0 E. haemoperoxidus 0174
1 E. haemoperoxidus 0175
2 E. haemoperoxidus 0176
0 E. haemoperoxidus 1251
1 E. haemoperoxidus 1252
2 E. haemoperoxidus 1253
3 E. haemoperoxidus 1254
4 E. haemoperoxidus 1255
0 E. haemoperoxidus 1538
1 E. haemoperoxidus 1539
2 E. haemoperoxidus 1540
3 E. haemoperoxidus 1541
4 E. haemoperoxidus 1542
5 E. haemoperoxidus 1543
-3 -4 E. haemoperoxidus 1823
-2 -3 E. haemoperoxidus 1824
-1 -2 E. haemoperoxidus 1825
0 -1 E. haemoperoxidus 1826
1 0 E. haemoperoxidus 1827
0 E. pernyi 0468
1 E. pernyi 0469
-5 E. pernyi 1663
-4 E. pernyi 1664
-3 E. pernyi 1665
-2 E. pernyi 1666
-1 E. pernyi 1667
0 E. pernyi 1668
B. Predicted toxins similar to epx4
Genomic context of the known epx4 toxin
Predicted toxin gene
Epx4
QueA-like
ATPase
Cry1Ac-like
PAP2
### of genes away from the toxin
Gene locus tag
-3 E. moraviensis 2667
-2 E. moraviensis 2668
-1 E. moraviensis 2669
0 E. moraviensis 2670
1 E. moraviensis 2671
2 E. moraviensis 2672
3 E. moraviensis 2673
4 E. moraviensis 2674
5 E. moraviensis 2675
0 -2 E. moraviensis 3236
1 -1 E. moraviensis 3237
2 0 E. moraviensis 3238
3 1 E. moraviensis 3239
4 2 E. moraviensis 3240
-4 E. haemoperoxidus 0351
-3 E. haemoperoxidus 0352
-2 E. haemoperoxidus 0353
-1 E. haemoperoxidus 0354
0 E. haemoperoxidus 0355
1 E. haemoperoxidus 0356
2 E. haemoperoxidus 0357
Structural alignment significance
Random (score < 0.4)
Borderline (0.4 < score < 0.5)
Nonrandom (0.5 < score < 0.6)
Significant (0.6 < score < 0.7)
Highly significant (score > 0.7)
Predicted toxin gene
Novel toxin gene predicted by only SAP

**Supplementary Fig. S7.** Detailed view of the genomic neighborhood of 11 likely novel toxin genes (red boxes under *epx* columns). Each row corresponds to a gene within the genomic neighborhood of the 11 predicted toxin genes. Genes are numbered according to their relative position within the operon in reference to the predicted toxin gene (column titled "# of genes away from the predicted toxin") and the corresponding gene locus tags, a 4-digit number given based on their location within the genome, are also listed in the column titled "Gene locus tag". A) Comparison to the known *epx1* neighborhood. B) Comparison to the known *epx4* neighborhood. The two operon diagrams at the top (adapted from Xiong et al., 2022) show the known genomic contexts of *epx1* and *exp4*, while the heatmaps below show protein structural alignment significance for proteins within this genomic context.

### References


Lebreton, F. *et al.* (2017). Tracing the enterococci from paleozoic origins to the hospital. *Cell*, **169**(5), 849–861.

Okuda, S. and Yoshizawa, A. C. (2010). Odb: a database for operon organizations, 2011 update. *Nucleic acids research*, **39**(suppl_1), D552–D555.

Parks, D. H. *et al.* (2021). Gtdb: an ongoing census of bacterial and archaeal diversity through a phylogenetically consistent, rank normalized and complete genome-based taxonomy. *Nucleic Acids Research*, **50**(D1), D785–D794.

Seemann, T. (2014). Prokka: rapid prokaryotic genome annotation. *Bioinformatics*, **30**(14), 2068–2069.

Xiong, X. *et al.* (2022). Emerging enterococcus pore-forming toxins with mhc/hla-i as receptors. *Cell*, **185**(7), 1157–1171.e22.

Zhou, N. *et al.* (2019). The CAFA challenge reports improved protein function prediction and new functional annotations for hundreds of genes through experimental screens. *Genome Biology*, **20**(1), 244.
